# A proteomic and phosphoproteomic comparison of mouse spermatogonial stem cells and progenitor spermatogonia

**DOI:** 10.64898/2026.07.15.738804

**Authors:** David A. Skerrett-Byrne, Brett Nixon, Adrián Sanz-Moreno, Filip Dámek, Patricia da Silva-Buttkus, Raffaele Teperino, Valérie Gailus-Durner, Helmut Fuchs, Martin Hrabě de Angelis, Jon M. Oatley, Connor Cason, Ilana R. Bernstein, Tessa Lord

## Abstract

In this manuscript, we used an *Id4-eGfp* mouse line to produce the first proteomic and phosphoproteomic database for mouse spermatogonial stem cells (SSCs) and progenitor spermatogonia. Our proteomic analyses identified 8,464 proteins in spermatogonia, superseding the depth of previously published spermatogonia datasets by >2000 proteins. While the comparison of SSCs and progenitors revealed few unique proteins (18 and 3, respectively), 532 proteins exhibited significantly different abundance (FC ± 1.5, *p*-value ≤ 0.05) between these sub-populations. Interestingly, in overlaying the proteome with transcriptomic data, correlation was poor (R^2^ = 0.236), re-iterating discordance between transcript and protein abundance in the testis. In our phosphoproteomic analyses, phosphosites were identified in 19.5% of proteins (3,604 total phosphosites). Unique protein phosphorylation was substantially more common than unique protein expression when comparing SSCs and progenitors, with 38 and 191 unique phosphosites identified, respectively, in addition to significant differences in abundance at 60 and 257 phosphosites. In identifying a wave of phosphorylation that accompanies the progenitor transition, we performed predictive analyses to identify three potential master kinases for follow up validation studies (PAK1, BUB1, ABL2). The inhibition of these kinases resulted in a significant reduction in the capacity for progenitor spermatogonia to differentiate, and caused elevated apoptosis and DNA damage. Finally, we explored the testis phenotype of 42 knockout mouse lines for proteins that were either differentially expressed (26) or differentially phosphorylated (15) in our dataset. Of these lines, 21 exhibited a testis phenotype in at least one of two biological replicates observed (severity score ≥ 1, compared to 0.15 for controls), with increased numbers of Sertoli-only tubules being evident in four of these lines. This manuscript provides a comprehensive roadmap to understand the multifaceted layers of regulation over fate decisions in undifferentiated spermatogonia. These data have been provided in an accessible platform via our ShinySpermatogoniaCells app to encourage future progress in the field.

## INTRODUCTION

The stem cells within the testis, or spermatogonial stem cells (SSCs), provide the basis for ongoing male fertility. The SSCs self-renew to maintain the undifferentiated reservoir, while simultaneously giving rise to progenitor spermatogonia that are poised to enter the differentiation pathway that terminates in mature sperm production. Intricate regulation over gene and protein expression, as well as protein/enzyme activity, is critically important for dictating fate decisions in stem cell populations^1,2^. However, a gap in knowledge exists surrounding the shift in protein abundance and post-translational modifications that occur in SSCs and progenitors at important developmental junctions.

In the previous decade, a wealth of transcriptomic data have been produced on SSCs and progenitor spermatogonia, primarily in rodent species, but also from human and primate testes^3–8^. This has largely been facilitated by the development of single-cell RNA sequencing (scRNA-seq) technologies that have revolutionised our ability to study transcript dynamics in rare cell populations. Although these datasets have been an invaluable tool driving significant progress in the field, the interpretation of transcriptomic findings can be limited by the known discordance between RNA and protein expression. Indeed, a recently published quantitative proteomic analysis revealed that, in the testis, over 50% of genes exhibit discordant expression, with a majority of these being enriched at the RNA level^9^. Beyond this, the activity of proteins is tightly regulated by post-translational modifications such as phosphorylation, with over 75% of the proteome estimated to be phosphorylated^10^. Thus, inferring function from levels of transcript expression alone misses several additional layers of regulation.

Despite the value in understanding the expression and modification of proteins within SSC and progenitor populations, very few proteomic analyses have been attempted, and to the authors knowledge, no phosphoproteomic databases exist for SSCs. Primarily, proteomic assessments have been performed on heterogeneous populations of undifferentiated spermatogonia (i.e. mixed populations of SSCs and progenitors)^11–14^, usually following a period of expansion in *in vitro* culture^12–14^. While useful, this approach overlooks the nuanced differences between these functionally distinct cell populations, and further, *in vitro* culture may alter protein composition from that of the original cell population *in vivo*^15^.

Roadblocks for producing proteomic and phosphoproteomic datasets for the SSC population have firstly been that these are an incredibly rare population of cells, comprising only 0.03% of the total cell population in the mouse testis^16^. Tied in with this, the sensitivity of proteomic and phosophoproteomic techniques has historically been much lower than that of transcriptomics, requiring large quantities of protein for in-depth analysis: a significant challenge when working with low cell numbers. The second roadblock has been the lack of markers that can facilitate isolation of pure populations of SSCs (reviewed in Lord and Oatley^17^), with existing methodologies achieving enrichment, but not purity of these stem cells. Excitingly, techniques have recently advanced such that phosphoproteomic approaches can now be carried out using as little as 25 µg of protein^18,19^. Alongside this, developments have been made in our ability to isolate populations of mouse SSC and progenitor spermatogonia. Specifically, the Oatley laboratory produced a transgenic reporter mouse line, the *Id4-eGfp* mouse^20^, which, using gold-standard limiting dilution spermatogonial transplantation techniques has been shown to facilitate the isolation of veritably pure SSC populations from the postnatal day (P)6-8 testis^21^. In this model, spermatogonia with the highest levels of *Id4* expression (ID4-eGFP^Bright^ cells) are the SSCs, while progenitor spermatogonia exhibit reduced *Id4* expression (i.e. are the ID4-eGFP^Dim^ cells). Conveniently, extensive transcriptomic analyses have already been conducted on these two populations, using both bulk RNAseq^21^ and single-cell RNAseq^3^, paving the way for complementary proteomic and phosphoproteomic analyses.

In this manuscript, we have used the *Id4-eGfp* mouse line to produce the first proteomic and phosphoproteomic database for mouse SSCs and progenitor spermatogonia and have overlaid these with existing RNAseq data to provide a comprehensive roadmap representing the transition from self-renewing to differentiation-poised cell. Further, we have conducted functional follow-up studies using kinase inhibitors and knockout mouse lines to highlight the important roles of differentially expressed or differentially phosphorylated proteins identified in our dataset. Ultimately, we have produced a comprehensive and accessible resource for the field via our Shiny Spermatogonia Cells app: https://reproproteomics.shinyapps.io/ShinySpermatogoniaCells/, with the hope that this resource will fuel future investigation into stem cell and reproductive biology alike.

## RESULTS

### Generating a proteomic database for SSC and progenitor spermatogonia

SSC and progenitor populations were isolated from *Id4-eGfp* transgenic mice at P6-8 using fluorescence activated cell sorting (FACS). SSCs were ID4-eGFP^Bright^ spermatogonia and progenitors were ID4-eGFP^Dim^, as reported previously^21^ and depicted in flow cytometry plots (Figure 1A). These enriched populations underwent the EasyPhos workflow^19,22^ to simultaneously profile their proteome and phosphoproteome (Figure 1A). Proteomic analysis returned a deep complex suite of proteins, with 8,461 and 8,446 proteins identified in the SSCs and progenitors respectively (Figure 1B, Table S1). Both cell types yielded an average of 15.2 unique peptides per protein, as well as 33.6% average protein coverage. This is in part due to the substantial overlap of protein IDs (99.8%), with 18 proteins uniquely detected in SSCs and a mere 3 in progenitors (Figure 1C). Amongst proteins unique to SSCs are those with known roles in chromatin regulation, SWI/SNF-related matrix-associated actin- dependent regulator of chromatin subfamily B member 1 (SMARCB1); Germ cell differentiation, transcription factor Ovo-like 2 (OVOL2); and piRNA biogenesis in germ cells, putative ATP-dependent RNA helicase TDRD12 (TDRD12) (Figure 1D). The three unique proteins to progenitors were cytoglobin (CYGB), SH3 and multiple ankyrin repeat domains protein 2 (SHANK2), and zinc finger CCCH domain-containing protein 14 (ZC3H14) (Figure 1D).

**Figure 1:**
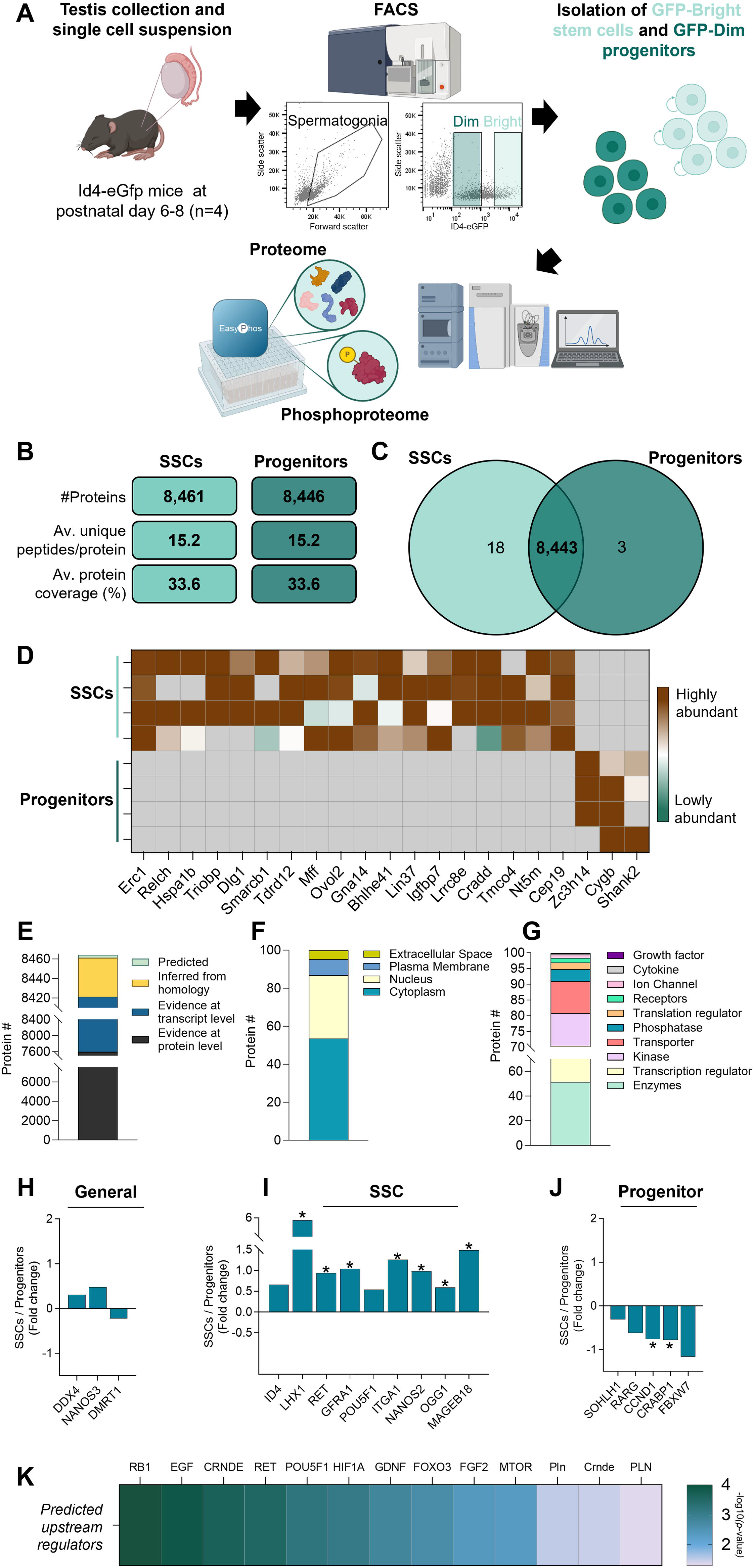
Proteomic analysis of mouse SSCs and progenitor spermatogonia. **A)** Pipeline by which SSCs and progenitor spermatogonia were isolated via FACS from P6-8 *Id4-eGfp* mice for EasyPhos analysis. **B)** Number of proteins identified, average unique peptides per protein, and average protein coverage (%) in SSC and progenitor populations. **C)** Venn diagram depicting the number of overlapping and unique proteins identified in SSCs and progenitors. **D)** Heatmap depicting expression abundance of the 18 proteins found to be uniquely expressed in SSCs, and 3 proteins uniquely expressed in progenitors, across n=4 independent biological replicates. (Brown = highly abundant, green = lowly abundant, grey = not detected). **E)** UniProt analysis depicting the number of identified proteins with annotated evidence at the protein level, evidence at the transcript level, inference from homology, or predicted based on the genome. **F)** Cellular localization of proteins, as predicted by Ingenuity Pathway Analysis (IPA). **G)** Protein type classification as predicted by IPA. **H)** Proteins known to be expressed by germ cells / spermatogonia exhibiting no significant difference in expression between SSCs and progenitors. Histogram depicts average fold-change of protein expression (SSCs / progenitors) across n = 4 independent biological replicates. **I)** Expression of proteins that have previously defined roles in SSC maintenance or self-renewal. Histogram depicts average fold-change of protein expression (SSCs / progenitors) across n = 4 biological replicates, * denotes significantly different at *p* ≤ 0.05. **J)** Expression of proteins that have previously defined roles in progenitor formation or in preparing progenitors for the differentiating transition. Histogram depicts average fold-change of protein expression (SSCs / progenitors) across n = 4 biological replicates, * denotes significantly different at *p* ≤ 0.05. **K)** Upstream regulators of spermatogonia predicted by an IPA assessment of differentially expressed proteins. Heatmap depicts the enrichment score (-log10 *p*-value).

Assessment of the full complement of proteins (8,464) with UniProt revealed 89.7% proteins have annotated evidence at the protein level (Figure 1E). The remaining 872 proteins currently possess evidence at the transcript level (829), are inferred from homology (40) or are predicted based on the genome (3). Given the high annotation completeness of the mouse proteome (99.8% completeness BUSCO score^23^ from UniProt), this 10.3% of low tier evidence suggests that these proteins may be spermatogonia and/or testis specific. Seeking to better understand the cellular localization and protein classifications, the proteome was next interrogated using Ingenuity Pathway Analysis (IPA). Cytoplasm and nucleus return the highest portions of the proteome, accounting for 53.5% and 33.3% respectively (Figure 1F, Table S1). The remaining 13.1% consisted of plasma membrane (8.5%) and extracellular space (4.6%). Protein type classification showcased that enzymes constituted the largest group, comprising 51.5% of the proteome (Figure 1G), followed by transcription regulators (19%), kinases (10.4%) and transporters (10.2%). Finally, the remaining 8.9% of the protein comprised of phosphatases, translation regulators, receptors, ion channels, cytokines and growth factors.

To further support the enrichment of spermatogonia sub-populations, the proteome was examined for established spermatogonia, SSC, and progenitor markers (Figure 1H–J). Classical spermatogonia markers ATP-dependent RNA helicase DDX4 (DDX4)^24^, Nanos homolog 3 (NANOS3)^25^ and Doublesex- and mab-3-related transcription factor 1 (DMRT1)^26^ were found to be in both populations with no significant difference in expression (Figure 1H). Several makers of SSCs were found to be significantly increased in the SSC population, including LIM/homeobox protein Lhx1 (LHX1)^27^, GDNF Family Receptor Alpha 1 (GFRA1)^28^ and Proto-oncogene tyrosine-protein kinase receptor Ret (RET)^29^ (Figure 1I). Contrastingly, proteins that promote progenitor formation (e.g. F-Box And WD Repeat Domain Containing 7 (FBXW7)^30^, and Cyclin D1 (CCND1)^31,32^) and those related to the retinoic acid response (e.g. Cellular retinoic acid-binding protein 1 (CRABP1)^33^), were found to be enriched in the progenitor population (Figure 1J). Moreover, the IPA upstream regulator function identified an additional 17 known regulators of SSCs, including Retinoblastoma 1 (RB1)^34^, Hypoxia Inducible Factor 1 (HIF1)^35,36^, Fibroblast Growth Factor 2 (FGF2)^37^ and Glial Cell Derived Neurotrophic Factor (GDNF)^38^ (Figure 1K). Collectively, these data confirm the successful enrichment of appropriate cell populations.

### Integrative Proteomic and Transcriptomic Analysis Reveals Key Functional Pathways Driving SSC Fate Decisions

To further contextualize the proteomic landscape uncovered here, we next integrated our data with a previously published RNA-seq dataset characterizing the same SSC and progenitor populations^21^. Unsurprisingly, >90% of the proteome overlapped with the reported transcriptome (Figure 2A). The 804 proteins not detected at the transcript level may be attributable to technological advancements and updated annotations introduced in the intervening years since the earlier study. Focusing on the changes observed between SSCs and progenitors, a Pearson correlation of the overlapping 7,660 proteins returned a low R^2^ of 0.236 (Figure 2B, Table S2). This is in line with previous studies reporting poor correlation between proteomics and RNA-seq datasets^39,40^. Despite this low correlation, 198 proteins were significantly altered (FC ± 1.5, *p*-value ≤ 0.05) in both datasets, with 174 sharing the same direction of change. Notably, Gametocyte-specific factor 1 (*Gtsf1*), DNA (cytosine-5)- methyltransferase 3B (*Dnmt3b*), and G1/S-specific cyclin-D1 (*Ccnd1*) displayed similar fold changes between the proteomic and transcriptomic analyses, underscoring their potential importance in the transition between SSCs and progenitors.

**Figure 2:**
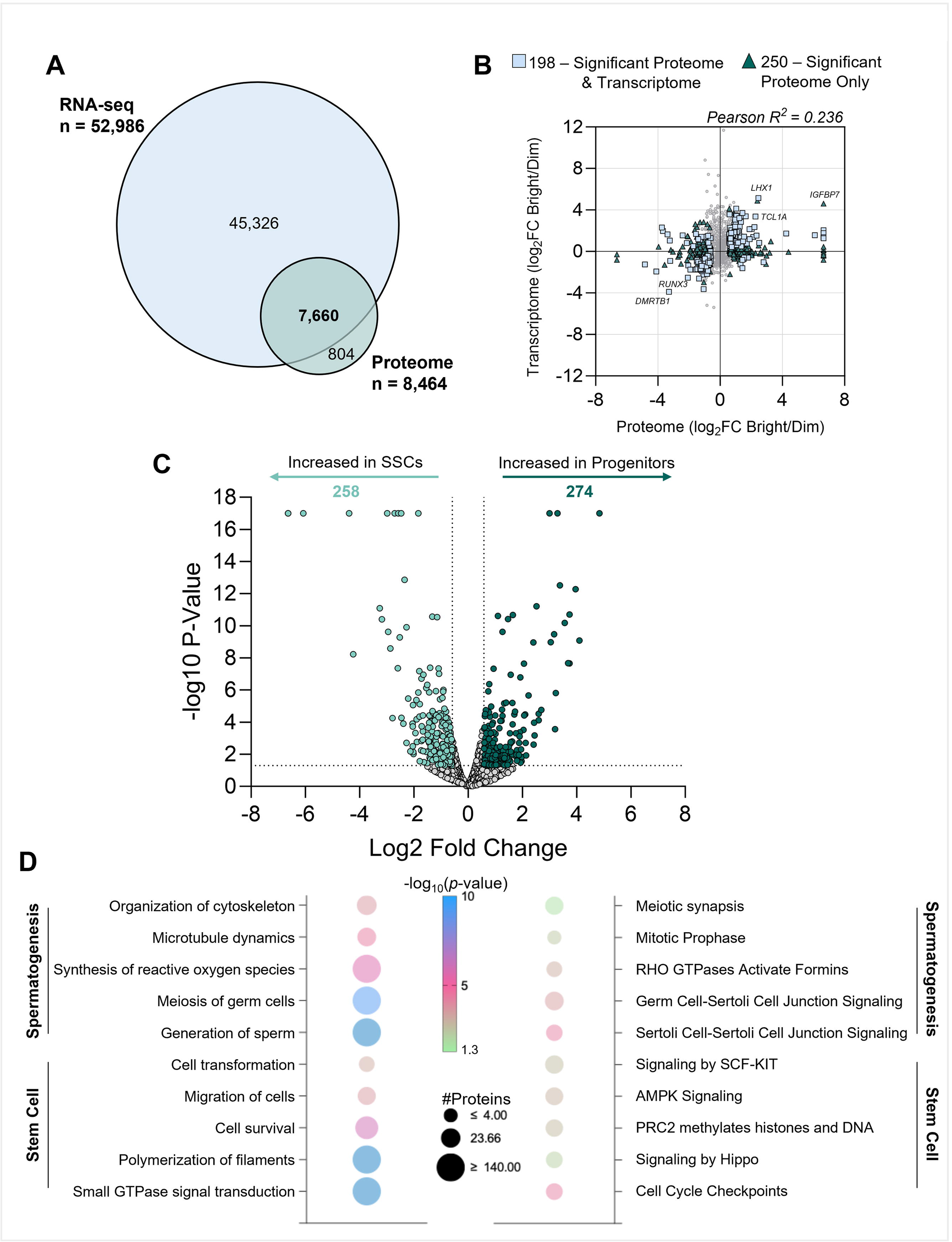
Assessing transcript and protein correlation and biological pathways associated with differentially expressed proteins. **A)** Venn diagram depicting the overlap between the proteomic dataset produced in this manuscript and a previously published RNAseq dataset capturing the same populations (ID4-eGFP Bright and Dim spermatogonia from postnatal mouse testes). **B)** Correlation of transcript and protein abundance between our proteomic dataset and previously published RNAseq data. Blue squares denote genes/proteins that have significantly different expression between SSCs and progenitors in both datasets, while green triangles represent factors that have significantly different expression in the proteome only. **C)** Volcano plot comparing protein expression in SSC versus progenitor spermatogonia. Points highlighted in green have a Log_2_ fold change exceeding ± 1.5, and *p*-value ≤ 0.05. **D)** Dot plot depicting ‘Diseases or Functions’ (left) and ‘Canonical Pathways’ (right) identified by IPA to be significantly enriched amongst differentially expressed proteins in our dataset. Dot size corresponds to the number of proteins in each category; dot colour corresponds to *p*-value. Highlighted pathways and functions have been categorized based on their relationship to Spermatogenesis or Stem Cell function.

Despite the extensive overlap in overall proteome composition between SSCs and progenitors (Figure 1C), volcano plot analysis highlighted substantial differences in protein abundance (Figure 2C, Table S1). In total, 532 proteins were significantly altered (FC ± 1.5, *p*-value ≤ 0.05) during the SSC-to-progenitor transition, with 274 increased and a reciprocal 258 decreased upon progenitor formation. Amongst the most significantly abundant proteins in progenitor cells were Activity-regulated cytoskeleton-associated protein (ARC, FC = 6.64), CD82 antigen (CD82, FC = 4.24) and Alkylated DNA repair protein alkB homolog 8 (ALKBH8, FC = 3.26). Conversely, the SSC population showed higher levels of Mitogen-activated protein kinase kinase kinase 9 (MAP3K9, FC = 4.84), Latent-transforming growth factor beta-binding protein 4 (LTBP4, FC = 3.74), and Doublesex- and mab-3-related transcription factor B1 (DMRTB1, FC = 3.29). To better understand the biological implications of these alterations, we employed IPA, enabling us to discern the key pathways and functional networks governing SSC fate decisions. Two major categories emerged from these analyses: those related to spermatogenesis (30 functions) and those linked to stem cell function (9 functions) (Figure 2D, Table S3). Spermatogenesis-related functions were highlighted by direct enrichment of the term ‘spermatogenesis’ and further supported by functions governing germ cell development and differentiation (‘generation of sperm’ and ‘gametogenesis’) (Table S3). Additionally, cytoskeletal and structural dynamics (‘organization of cytoskeleton’ and ‘microtubule dynamics’) and metabolic regulation (‘synthesis of reactive oxygen species’) reinforced the involvement of processes essential for producing and maintaining healthy male germ cells. Functions pertinent to stem cell biology included those essential for maintaining a proper niche environment (‘cell transformation’ and ‘cell survival’), guiding stem cell self-renewal and differentiation (‘small GTPase mediated signaling transduction’), and facilitating the cellular dynamics necessary for stem cell development and repair (‘migration of cells’ and ‘polymerisation of filaments’).

In complement to the functional annotation, canonical pathway analysis echoed the two overarching categories, those predominantly engaged in spermatogenesis and those underpinning stem cell related programs (Figure 2E, Table S3). Spermatogenesis-associated pathways included ‘Sertoli-Cell–Sertoli-Cell Junction Signaling’ and the complementary ‘Germ Cell–Sertoli Cell Junction Signaling’, together underscoring coordinated adhesion and paracrine support required by spermatogonia. Stem cell-oriented pathways converged on complementary mechanisms that safeguard SSC self-renewal while permitting timely commitment to differentiation. Genomic vigilance was underscored by ‘Cell Cycle Checkpoints’, which enforce G1/S and G2/M surveillance to avert mutation accumulation during rapid proliferative bursts^41^. Metabolic poise emerged via ‘AMPK Signaling’, a key energy sensor that keeps quiescent cells in a low-ATP, high-autophagy state until external cues signal entry into the cell cycle^42^. Long-term epigenetic memory was highlighted by ‘PRC2 Methylates Histones and DNA’, pointing to Polycomb-mediated H3K27 trimethylation that maintains transcriptional repression of differentiation genes^43^. Context-dependent growth control was illustrated by ‘HIPPO Signaling’, whose YAP/TAZ effectors fine-tune the balance between proliferation and apoptosis within stem cell compartments^44,45^). Finally, ‘RET signaling’ and ‘Signaling by SCF–KIT’ anchored the importance of paracrine support from Sertoli cells^46,47^, providing stimuli to drive self-renewal and differentiation, respectively. Taken together, these findings offer mechanistic insights into how shifts in the proteome may guide SSC fate decisions and shape the progenitor state.

### Characterization of the phosphoproteomic changes underpinning SSC differentiation

Simultaneous with our proteomic analysis, we enriched for corresponding phosphopeptides and sequenced 3,633 and 3,790 phosphosites from SSCs and progenitors respectively (Figure 3A, Table S4), with phosphosites ultimately being detected in 19.5% of proteins in the proteome (Figure 3B, Table S1). When comparing at the peptide level, 38 phosphopeptides were uniquely detected in the SSC population, while a comparatively substantial 191 phosphopeptides were detected in progenitor spermatogonia (Figure 3A). In SSCs, proteins mapping to these uniquely phosphorylated sites included CREB binding protein (CREBBP), a member of the CREB complex that we have previously shown is important for SSC self-renewal^48^. Proteins uniquely phosphorylated in progenitors included Testis Expressed 14, Intercellular Bridge Forming Factor (TEX14), an essential protein for intercellular bridge formation of progenitor spermatogonia^49^, and RPTOR Independent Companion Of MTOR Complex 2 (RICTOR), the expression of which is essential for the transition of progenitor spermatogonia into a differentiating state^50^.

**Figure 3:**
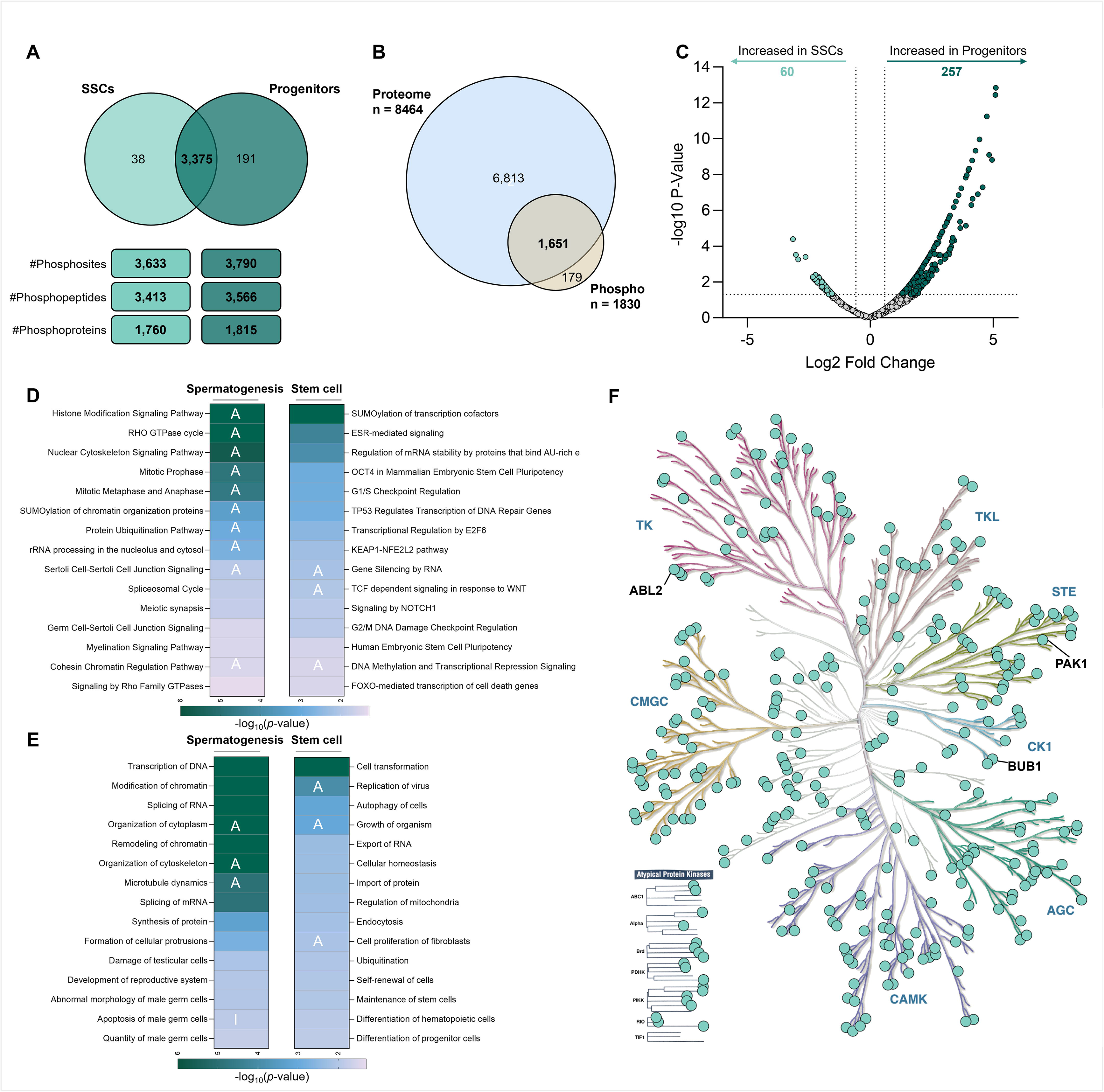
Phosphoproteomic analysis of mouse SSCs and progenitor spermatogonia. **A)** (Upper) Venn diagram depicting the number of overlapping and unique phosphopeptides identified in SSCs and progenitors. (Lower) Number of phosphosites, phosphopeptides, and phosphoproteins identified in SSCs and progenitors. **B)** Venn diagram highlighting the overlap between phosphoproteins and the total proteome, identified across SSC and progenitor populations combined. **C)** Volcano plot comparing phosphopeptides in SSC versus progenitor spermatogonia. Points highlighted in green have a Log2 fold change exceeding ± 1.5, and *p*-value ≤ 0.05. **D)** Heat map depicting ‘Canonical Pathways’ identified by IPA to be significantly enriched amongst differentially phosphorylated proteins in our dataset. Colour corresponds to enrichment scores (- log_10_*p*-value). ‘A’ (activation) denotes a Z score ≥2. Highlighted pathways have been categorized based on their relationship to Spermatogenesis (left) or Stem Cell function (right). **E)** Heat map depicting ‘Diseases or Functions’ identified by IPA to be significantly enriched amongst differentially phosphorylated proteins in our dataset. Colour corresponds to enrichment scores (- log_10_*p*-value). ‘A’ (activation) denotes a Z score ≥ 2, ‘I’ (inhibition) denotes a Z score ≤ -2. Highlighted pathways have been categorized based on their relationship to Spermatogenesis (left) or Stem Cell function (right). **F)** Kinome dendrogram (via KinMap) displays the distribution of the 328 identified kinase candidates potentially governing spermatogonia differentiation.

In considering phosphopeptides that were identified in both SSC and progenitor populations, further substantial differences were identified, with 60 phosphopeptides being significantly over-represented in the SSC population (FC ± 1.5, *p*-value ≤ 0.05) and 257 over-represented in progenitors (Figure 3C, Table S4). Proteins that were found to be differentially phosphorylated that have well-defined roles in dictating spermatogonia fate included ID4 itself, for which Ser5 phosphorylation was significantly overrepresented in SSCs. Given that Ser5 phosphorylation in the closely related protein, ID2, has been found to lock cells into a stem cell transcriptional state^51^, one can posit that that differential phosphorylation of ID4 could act as an additional layer of regulation to sustain SSC self-renewal. Further to ID4, EPHA2^52^, and members of the NuRD complex (H3-3A, DAXX, CHD4) that we have previously characterised as having a key role in SSC maintenance^53^, were also identified as being differentially phosphorylated in SSCs versus progenitors.

In extending our systems-level interrogation to the phosphoproteome, IPA analysis once again revealed two dominant, phosphorylation-sensitive networks; spermatogenesis and stem cell regulation (Table S5). Spermatogenesis-associated pathways encompassed chromatin and cytoskeletal remodeling events critical for germ-cell maturation. Prominent among these were activation of ‘Histone Modification Signaling’, highlighting the dynamic epigenetic landscape required to pre-empt differentiation^54^, and activations of the ‘RHO GTPase Cycle’, which orchestrates actin filament rearrangements essential for the cell cycle and cytokinesis ^55^. Also captured was ‘Germ Cell-Sertoli Cell Junction Signaling’, again highlighting the important role of Sertoli cells in dictating spermatogonia fate. Finally, activation of the ‘Spliceosomal Cycle’ may point to the extensive post-transcriptional regulation that is required for spermatogonial differentiation^56^ (Table S5). Stem cell pathways converged on mechanisms that balance self-renewal with genome integrity (Figure 3D). The role of ‘OCT4 in Mammalian Embryonic Stem Cell Pluripotency’ underscored core transcriptional circuitry sustaining an undifferentiated state^57^, whereas ‘Cell-Cycle Checkpoints (G1/S and G2/M)’ reflected heightened surveillance to preserve genomic stability in proliferating SSCs^58^. Redox-sensitive control was evident through the ‘KEAP1- NFE2L2 Pathway’, potentially pointing to an adaptive antioxidant response within the stem cell niche^59^. Developmental cell-fate cues emerged via ‘NOTCH1 Signaling’^60^, and long-term epigenetic memory was implied by activations of ‘DNA Methylation and Transcriptional Repression Signaling’, collectively delineating a multifaceted regulatory framework guiding SSC maintenance and commitment (Table S5).

Our phosphoproteomic dataset highlights the central role of kinase signaling in sculpting fate decisions in undifferentiated spermatogonia, in particular suggesting a prominent role for kinases in progenitor formation and/or the preparation of progenitors for the differentiating transition. To provide further biological insight, we endeavoured to predict key kinases that promote differential phosphorylation in SSCs versus progenitors. Using an *in silico* pipeline developed previously^19,61^ that uses IPA, PhosphoSitePlus^62^, and proteome data published in this manuscript, we identified 328 kinases spanning 8 major kinase families (Figure 3F, Table S6). Importantly, candidates predicted by this pipeline included numerous kinases with well-established roles in regulating spermatogonia function including RET^29^, EPHA2^52^, AKT1^27^ and MAP2K1^37^, highlighting the validity of this pipeline. Beyond this, an abundance of novel candidates were identified (Table S6), which provide fuel for future functional and validation studies.

### Master kinases PAK1, BUB1 and ABL2 protect spermatogonia integrity and prepare progenitors for differentiation

In order to complement the identification of putative master kinases predicted to play key roles in undifferentiated spermatogonia (Figure 3F), we performed validation experiments for three targets for which pharmacological inhibitors already exist: PAK1, BUB1 and ABL2. The inhibitors NVS-PAK1-1, BAY-1816032 and Asciminib, were used to treat undifferentiated spermatogonia in culture at concentrations of 2.5nM, 25nM and 25nM, respectively (24h incubation), and a fourth treatment group was created in which the three kinases were inhibited simultaneously (“Trio”) (Figure 4). Concentrations were selected based on the results of dose response experiments (Figure S1) and previously published IC50 values^63–65^. Using the percentage of ID4-eGFP^Bright^ and ID4-eGFP^Dim^ cells in each treatment as a proxy for SSC and progenitor abundance (as described previously^48^), no change the ratio of SSCs and progenitors within the undifferentiated spermatogonia population could be identified following kinase inhibition (Figure 4A). However, in assessing the percentage of differentiating spermatogonia in the population in response to a 6h DMSO (vehicle) or retinoic acid (0.25µM) treatment (Figure 4B), a significant reduction in the percentage of cKIT+ spermatogonia was observed across all treatments (p<0.05). As we and others have reported previously, the direct application of retinoic acid *in vitro* drives differentiation indiscriminately in all spermatogonia (i.e. ID4-eGFP^Bright^ and ID4-eGFP^Dim^: demonstrated in flow cytometry plots in Figure 4B), in contrast to the nuanced action on progenitors specifically that occurs within the niche environment *in vivo*^66,67^. Regardless, these experiments suggest a co-operative role for PAK1, BUB1 and ABL2 in the preparation of progenitor spermatogonia for differentiation. This aligns with the identification of ‘Differentiation of progenitor cells’ as a significantly enriched pathway amongst proteins with differential phosphorylation in Figure 3E.

**Figure 4:**
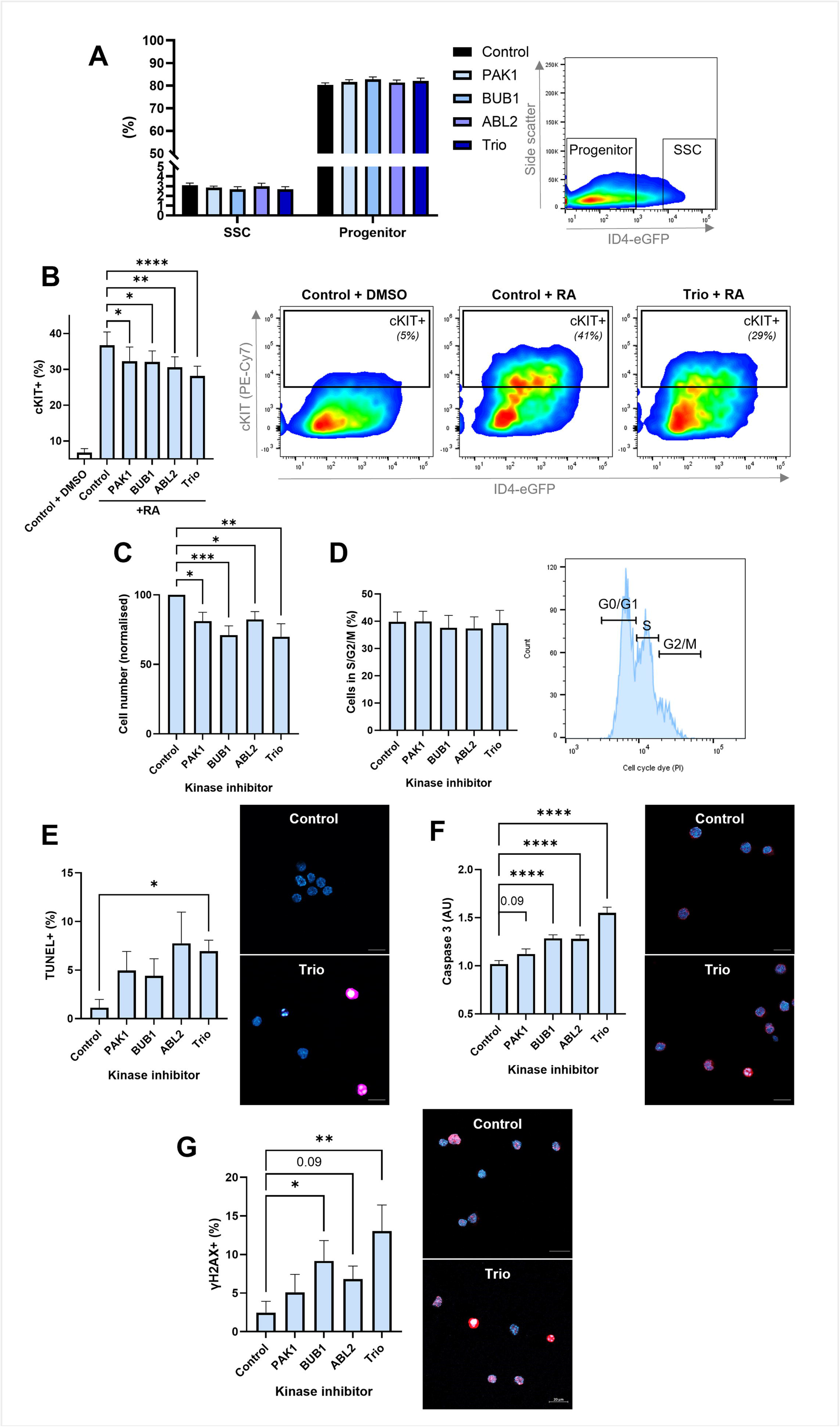
Inhibition of predicted master kinases in undifferentiated spermatogonia in culture. **A)** Percentage of ID4-eGFP^Bright^ SSCs and ID4-eGFP^Dim^ progenitors in primary cultures of undifferentiated spermatogonia following pharmacological inhibition of the kinases PAK1, BUB1, ABL2, or all three concurrently (Trio’), as compared to a DMSO vehicle control. Left: Histogram depicts mean ± SEM for n = 4 independent biological replicates. Right: Representative flow cytometry plot showing gating parameters. **B)** Percentage of cKIT+ differentiating spermatogonia in primary cultures of undifferentiated spermatogonia treated with kinase inhibitors for PAK1, BUB1, ABL2, or Trio, followed by retinoic acid (RA). Control cells treated with vehicle (DMSO) are provided as a comparison. Left: Histogram depicts mean ± SEM for n = 5 independent biological replicates. * denotes significantly different at p < 0.05, **** denotes significantly different at p < 0.0001. Right: Representative flow cytometry plots for control samples treated with DMSO or RA, and Trio samples treated with RA. **C)** Number of undifferentiated spermatogonia in culture (normalised to control) following treatment with kinase inhibitors. Histogram depicts mean ± SEM for n = 10 independent biological replicates. * denotes significantly different at p < 0.05, ** denotes significantly different at p < 0.01, *** denotes significantly different at p < 0.001. **D)** Cell cycle analysis depicting the percentage of undifferentiated spermatogonia in primary culture in S/G2/M phase following treatment with kinase inhibitors for PAK1, BUB1, ABL2, or Trio. Left: Histogram depicts mean ± SEM for n = 6 independent biological replicates. Right: Representative flow cytometry plot showing gating parameters. **E)** Percentage of TUNEL positive undifferentiated spermatogonia following treatment of primary cultures with kinase inhibitors for PAK1, BUB1, ABL2, or Trio, as compared to a vehicle (DMSO) treated control. Left: Histogram depicts mean ± SEM for n = 4 independent biological replicates. * denotes significantly different at p < 0.05. Right: Representative images for control and Trio treated cells (additional images provided in Figure S2). Blue staining is DAPI, purple staining is TUNEL. Scale bar = 20 µm. **F)** Fluorescence intensity analysis (normalised to control) of cleaved Caspase 3 in undifferentiated spermatogonia following treatment of primary cultures with kinase inhibitors for PAK1, BUB1, ABL2, or Trio. Left: Histogram depicts mean ± SEM for n = 3 independent biological replicates. **** denotes significantly different at p < 0.0001. Right: Representative images for control and Trio treated cells (additional images provided in Figure S2). Blue staining is DAPI, red staining is Caspase 3. Scale bar = 20 µm. **G)** Percentage of γH2AX positive undifferentiated spermatogonia following treatment of primary cultures with kinase inhibitors for PAK1, BUB1, ABL2, or Trio, as compared to a vehicle (DMSO) treated control. Left: Histogram depicts mean ± SEM for n = 4 independent biological replicates. * denotes significantly different at p < 0.05, ** denotes significantly different at p < 0.01. Right: Representative images for control and Trio treated cells (additional images provided in Figure S2). Blue staining is DAPI, red staining is γH2AX. Scale bar = 20 µm.

Inhibition of PAK1, BUB1, ABL2 and ‘Trio’ also resulted in a significant reduction in overall cell number in the undifferentiated spermatogonia population (Figure 4C, p<0.01). To identify the underlying cause, we firstly explored cell cycle activity (Figure 4D) however did not identify any change in the percentage of cells in S/G2/M phase. Contrastingly, when we explored markers of apoptosis, a significant increase in the percentage of TUNEL+ cells was apparent in the ‘Trio’ treatment (p<0.05, Figure 4E, S2), as was a significant elevation in Caspase 3 staining in response to BUB1, ABL2 and ‘Trio’ inhibitors (p<0.001, Figure 4F, S2), approaching significance in response to PAK1 inhibition (p=0.09). This aligns with the identification of ‘Apoptosis of male germ cells’ and ‘Quantity of male germ cells’ as significantly enriched pathways amongst proteins with differential phosphorylation (Figure 3E). Finally, we assessed the percentage of spermatogonia harbouring DNA damage (double strand breaks) using the marker γH2AX. Here we identified a significantly increased percentage of γH2AX+ spermatogonia in response to BUB1 and Trio inhibitors (p<0.0001, Figure 4G, S2), approaching significance in response to ABL2 inhibition (p=0.09), aligning with the enriched pathways ‘Damage of testicular cells’, ‘TP53 regulates transcription of DNA repair genes’ and ‘G2/M DNA damage checkpoint regulation’ (Figure 3E).

These mechanistic studies give credence to importance of the differentially phosphorylated proteins identified in our study, and demonstrate how dynamic changes in protein phosphorylation accompanying the SSC-to-progenitor transition are key for protecting the integrity of these cells while also preparing them for the significant remodeling that ensues upon differentiation / reception of the retinoic acid pulse.

### Testis phenotypes following knockout of genes encoding differentially expressed and phosphorylated proteins

Finally, to provide further insight into the role of differentially expressed and phosphorylated proteins identified in our study, we examined testes and epididymides from 42 knockout mouse lines generated at the German Mouse Clinic^68–70^. 26 of the genetic knockout lines were selected based on their protein product being significantly enriched either in SSCs or progenitors (Figure 5A, left), while 15 were chosen based on significantly different protein phosphorylation (Figure 5B, left), and 1 was picked based on a significant differences in protein expression and phosphorylation (VGLL4, Figure 5A, B, left).

**Figure 5:**
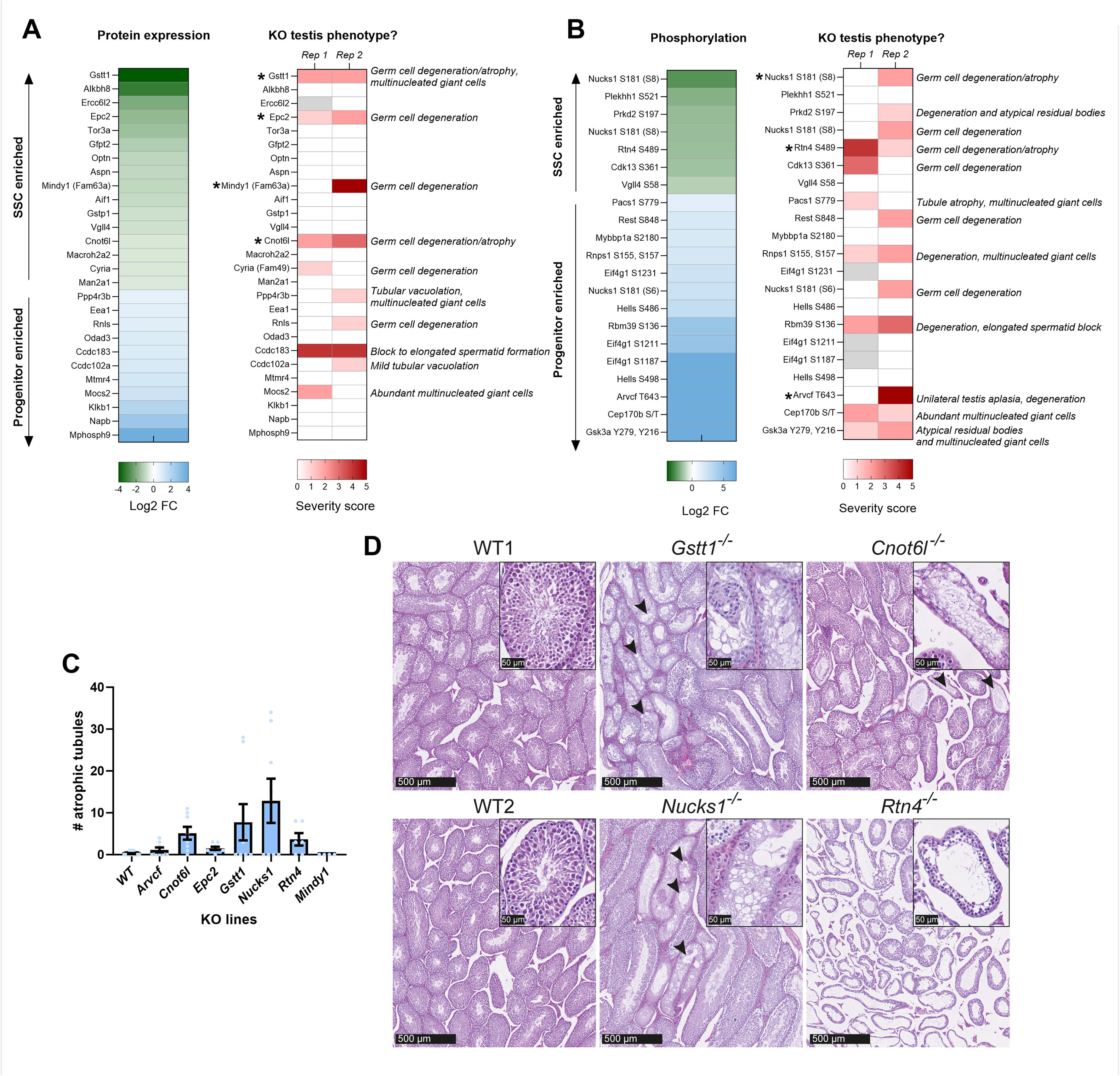
Assessing testis phenotypes in knockout mouse lines for proteins and phosphoproteins highlighted in our analysis. **A)** Testis phenotype severity in knockout mouse lines for genes whose protein products were differentially expressed in SSCs versus progenitors. Left: Heatmap showing the Log_2_ fold change of proteins that were selected for follow up analysis. Green = proteins significantly enriched in SSCs, blue = proteins significantly enriched in progenitors. Right: Heatmap showing the testis phenotype severity score for each knockout line (n = 2). A score of 0 (white) indicates no detectable abnormalities, and scores 1, 2, 3, 4 and 5 (represented by scale of light to dark red) correspond to mild, moderate, marked, marked-to-severe and severe changes, respectively. **B)** Testis phenotype severity in knockout mouse lines for genes whose protein products were differentially phosphorylated in SSCs versus progenitors. Left: Heatmap showing the Log2 foldchange of phosphoproteins that were selected for follow up analysis. Green = phosphoproteins significantly enriched in SSCs, blue = phosphoproteins significantly enriched in progenitors. Right: Heatmap showing the testis phenotype severity score for each knockout line (n = 2). A score of 0 (white) indicates no detectable abnormalities, and scores 1, 2, 3, 4 and 5 (represented by scale of light to dark red) correspond to mild, moderate, marked, marked-to-severe and severe changes, respectively. **C)** Comparison of the number of Sertoli cell-only / atrophic tubules across in knockout mouse lines as compared to age matched controls. Histograms depict mean ± SEM. Knockout lines are n = 2 biological replicates and n = 6-8 technical replicates (individual testis sections). For wild type comparison, analysis was performed across n = 6 biological replicates and n = 24 technical replicates. **D)** Photomicrographs of hematoxylin and eosin-stained testes from 16-week-old wild-type (WT) control and *Gstt1-/-, Cnot6l-/-, Nucks1-/-* and *Rtn4-/-* mice. Sections from the mutant lines show clusters of Sertoli cell-only tubules. Insets show higher magnification views of cross-sections of seminiferous tubules. Additional images are provided in Figure S3.

For each line, formalin-fixed and paraffin-embedded testes and epididymides from two knockout mice and two genetic-background-matched controls at 16 weeks of age were analysed by histopathological evaluation (Table S7 and Figure S3). H&E-stained sections were annotated using a severity score (see Methods) developed by the IMPC Infertility working group (Figure 5A, B, right). A score of 0 represents no pathological abnormalities, whereas a score of 5 denotes severe histological alterations throughout the section likely to be associated with clinical manifestations. Testes from 82 control animals were assessed, with an average severity score of 0.15 (Table S7, Figure 5A). Of 27 knockout lines selected based on significant differences in protein abundance between SSCs and progenitors, 10 displayed a testis phenotype with severity score of ≥1 in at least one replicate (Figure 5A). Similarly, of 16 knockout lines selected based on differential protein phosphorylation in SSCs versus progenitors, 11 displayed a testis phenotype with severity score ≥1 in at least one replicate (Figure 5B).

We elected to do follow up analysis on seven knockout lines in which degeneration and depletion of early germ cell stages was evident in a subset of seminiferous tubules^71^, which we reasoned could be related to spermatogonia dysfunction. These lines were knockouts of the following genes: Glutathione S-Transferase Theta 1 [*Gstt1*]; Enhancer Of Polycomb Homolog 2 [*Epc2*]; MINDY Lysine 48 Deubiquitinase 1 [*Mindy1*]; and CCR4-NOT

Transcription Complex Subunit 6 Like [*Cnot6l*]) (Figure 5A, asterisks); Nuclear Casein Kinase And Cyclin Dependent Kinase Substrate 1 [*Nucks1*]; Reticulon 4 [*Rtn4*]; and ARVCF Delta Catenin Family Member [*Arvcf*] (Figure 5B, asterisks). In conducting a quantitative assessment of atrophic tubules in the testes from each line, it could be appreciated that despite variability between animals, the mean number of Sertoli cell-only tubules identified in *Cnot6l* (5.1 ± 1.5)*, Gstt1* (7.8 ± 4.3)*, Nucks1* (12.9 ± 5.3) and *Rtn4* (3.7 ± 1.5) knockout animals was noticeably elevated from that of wild type (0.3 ± 0.1) (Figure 5C, D, Figure S3, noting that statistical tests were not performed given that only two biological replicates were available for knockout lines).To further extrapolate this, we performed immunostaining to identify any disruption to the number of undifferentiated spermatogonia per tubule (LIN28+ cells, Figure S4A,C), or to the number of tubules containing differentiating spermatogonia (STRA8+, Figure S4B, D), i.e. the number of tubules in stage VII-VIII of the cycle of the seminiferous epithelium. Although only modest differences in the number of LIN28+ cells per tubule were observed (e.g. 5.0 ± 0.3 in the *Nucks1* knockout versus 4.4 ± 0.2 in wild type), trends towards a reduced number of tubules containing differentiating spermatogonia were evident, particularly in *Cnot6l* knockout (28.8 ± 0.4) and *Nucks1* knockout (27.9 ± 2.6) lines, as compared to wild type (36.9 ± 2.6), potentially reflecting impairment in the undifferentiated to differentiating transition. Although future experiments are required with additional replicates to identify statistical significance, increased rates of phenotypic change in a number of these knockout lines again validates the importance of the candidates highlighted in this study for spermatogenesis and spermatogonial function specifically.

## DISCUSSION

Understanding of how fate decisions are regulated in undifferentiated spermatogonia has potential applications that span from improved diagnosis of male idiopathic infertility to the development of novel fertility treatments. For instance, the development of *in vitro* pipelines that elicit nuanced control over SSC expansion and differentiation could have direct application for reversal of infertility in survivors of paediatric cancers, as well as in biobanking pipelines aimed at maintaining genetic diversity in vulnerable wildlife species (reviewed in ^72^ and ^73^). While an abundance of transcriptomic databases are now available for the study of gene expression profiles that are associated with self-renewal of SSCs and the transition to a differentiation-primed progenitor state (e.g. ^21,74^), this limited focus to the transcript level misses important layers of information on protein expression and regulation of activity by post-translational modifications. In this study, we have produced a new, comprehensive database for the field that captures the proteome and phosphoproteome of mouse SSCs and progenitor spermatogonia, which we have made publicly accessible via an interactive Shinyapp: “ShinySpermatogoniaCells” (https://reproproteomics.shinyapps.io/ShinySpermatogoniaCells/ ). Our App enables on-the-fly queries for protein and gene expression, protein phosphorylation, and pathways enriched in the SSC-to-progenitor transition. This provides a valuable resource for fertility researchers and stem-cell biologists alike, and will serve as a benchmark for future single-cell proteomics analyses and cross-species comparisons (e.g., livestock, marsupials).

Our integrative analysis reinforces the substantial discordance between transcript and protein abundance,^40,75^ highlighting widespread post-transcriptional and post-translational control within the germline. Transcriptomic measurements represent a temporal snapshot of mRNA abundance, which does not necessarily reflect protein stability, translational efficiency, or regulated protein turnover. Consistent with this, proteins such as the DNA repair factor RecQ-mediated genome instability protein 2 (RMI2), the vesicular trafficking regulator ADP-ribosylation factor 2 (ARF2), and Protein FAM184B exhibited significantly high protein expression despite discordant transcript-level detection (Figure 2B, Table S2). Notably, several proteins that were differentially expressed at the protein level were not detected in the corresponding RNA-seq dataset, such as dynein axonemal heavy chains 12 and 17 (Table S2). Despite this absence at the transcript level, global knockout of genes such as *Nucks1* and *Cnot6l* resulted in marked testicular pathology, including germ cell degeneration/atrophy (Figure 5). Collectively, these findings provide functional evidence that reliance on transcriptomic profiling alone would have overlooked critical regulators of spermatogonial integrity and underscore the necessity of integrating proteomic and phosphoproteomic approaches when dissecting germ-cell biology.

In comparing the phosphoprotein landscape of SSCs and progenitor spermatogonia, we have identified significant changes in protein phosphorylation in the absence of corresponding changes in protein and/or transcript expression, highlighting kinase-substrate rewiring as a rapid, reversible mechanism for dictating fate decisions in spermatogonia. These differentially phosphorylated candidates (Figure 3C, Table S4) provide an untapped resource of novel factors to explore in future studies and explain some prior functional observations in investigations into SSC regulation. For example, high levels of serine phosphorylation were identified in CREBBP in SSCs that could no longer be detected upon the progenitor transition. Given that phosphorylation of CREBBP at serine residues has been linked to modulation of its function as a transcriptional co-activator^76^, these findings complement our previous research in which knockdown of *Crebbp* (and other members of the CREB complex) in undifferentiated spermatogonia was shown to impair SSC function specifically^48^, despite a lack of differential *Crebbp* expression between SSCs and progenitors at the transcript and protein level. Further, our previous study suggested that the CREB complex is important for directing histone acetylation in SSCs to influence expression of key self-renewal genes^48^. Thus, the differential phosphorylation of CREBBP is a prime example of how simple changes in protein phosphorylation can initiate a cascade of epigenetic, gene and protein expression alterations that dictate SSC fate decisions.

A similar principle is illustrated by additional candidates within our dataset, including NUCKS1. At a protein level, NUCKS1 exhibited no significant changes in overall abundance between SSCs and progenitors, however, displayed differential phosphorylation across the transition. Importantly, global deletion of *Nucks1* resulted in testicular pathology, with mutants exhibiting tubular atrophy (Sertoli cells only), and a decreased number of tubules with STRA8+ differentiating spermatogonia (Figure 5, S4). These findings further support the concept that phosphorylation-dependent modulation of protein function, rather than changes in steady-state abundance, can have profound consequences for spermatogonial integrity.

To further explore the notion that targeting phospho-nodes could modulate spermatogonia behaviour without altering gene expression, we used our previously established bioinformatic pipeline^19,61^ to predict master kinases based on changes in phosphorylation upon the SSC-to-progenitor transition (Figure 3F). Commercially available pharmacological inhibitors were used to consolidate a role for three of the predicted master kinases (PAK1, BUB1, and ABL2) in the coordinated priming of progenitor spermatogonia for differentiation and the protection of cell integrity. In alignment with predictions by IPA pathway analysis (Figure 3D, E), inhibition of this triad of kinases resulted in significantly elevated levels of apoptosis (Figure 4E, F) and DNA damage (Figure 4G) in undifferentiated spermatogonia in culture. This is cohesive with previously reported roles for these kinases. For instance, knockdown of *Pak1* in a human spermatogonia cell line xenografted into mice caused significantly elevated levels of apoptosis^77^, while knockdown in a HCT-116 cancer cell line caused significantly increased vulnerability to DNA double strand break formation in response to ionizing radiation^78^. Knockdown of *Bub1* has similarly been shown to instigate increased levels of DNA double strand breaks in HeLa cells because of impaired DNA damage repair, accompanied by a significant reduction in cell survival^79^, and impaired activity of ABL kinases has a well appreciated impact on cell survival and DNA integrity based on their interest as a target for leukemia treatment^80,81^. In our analyses, inhibition of PAK1, BUB1 and ABL2 also resulted in a significant reduction in the differentiation capacity of spermatogonia (Figure 4B). In the case of BUB1, this is likely intertwined with its ability to phosphorylate STAT3^82^, which has been shown to upregulate expression of *Neurog3* in progenitor spermatogonia to facilitate the differentiating transition^83^. ABL2 is predicted to be a component of the cellular response to retinoic acid, and the co-administration of retinoids with and ABL kinase inhibitor in a BCR-ABL1 leukemia cell line potentiated inhibitor activity, further suggesting co-regulation^84^. Overall, these pharmacological inhibition experiments underscore the non-redundant roles for PAK1, BUB1, and ABL2 in modulating spermatogonia behaviour, and provide entry points for fertility therapeutics, particularly in targeting DNA damage and apoptosis pathways. Additionally, these experiments highlight the enrichment in activity of cell-cycle checkpoints and DNA repair modules caused by the wave of phosphorylation accompanying the progenitor transition, demonstrating the stringent genomic safeguarding carried out in spermatogonia to preserve fertility across generations.

A prominent theme emerging from both our proteomic and phosphoproteomic analyses is the proactive management of oxidative stress within SSCs. Pathway interrogation showcased enrichment of the KEAP1-NRF2 axis alongside enrichment of detoxification enzymes (Figure 3D), particularly members of the glutathione S-transferase (GST) family. Such up-regulation suggests that SSCs maintain an anticipatory ROS-buffering state rather than responding passively to oxidative insult. Among these, GSTT1 was significantly enriched at the protein level in SSCs and its deletion resulted in germ cell degeneration and multinucleated giant cells, accompanied by hypospermia in the epididymis (Table S7). This phenotype reinforces the functional importance of antioxidant defence mechanisms in preserving early germ-cell integrity.^71^ Beyond detoxification, our dataset also revealed enrichment and predicted activation of base excision repair (BER) pathways in SSCs, which safeguard the genome against oxidative DNA lesions. This aligns with previous reports demonstrating high levels of BER and OGG1 activity in SSCs^85^ and supports the concept that redox buffering and DNA repair operate in concert to protect the long-term genomic stability of the stem-cell compartment. Collectively, these findings position antioxidant capacity as a strong feature of SSC identity, and not just a stress response pathway^86,87^. In a lineage tasked with transmitting genetic information across generations, stringent oxidative-stress management is likely a foundational characteristic.

In parallel with enhanced antioxidant defences, pathway interrogation revealed enrichment of metabolic and bioenergetic programs, including glucose metabolism, acetyl-CoA biosynthesis, and insulin receptor signalling, together with activation of energy-sensing pathways such as AMPK. These signatures suggest coordinated metabolic remodelling during the SSC-to-progenitor transition rather than metabolic stress. Although increased metabolic flux is often associated with elevated reactive oxygen species, the concurrent enrichment of KEAP1-NFE2L2 signalling and cytoprotective pathways (e.g., HMOX1), alongside DNA damage surveillance modules including ATM signalling, TP53-regulated repair, and base excision repair, supports a model of controlled mitochondrial and metabolic adaptation. We have previously demonstrated that spermatogonial differentiation is accompanied by shifts in mitochondrial function and bioenergetic output.^87^ The present data extend this framework by indicating that early fate transitions are underpinned by enhanced metabolic capacity, likely supporting increased biosynthetic and proliferative demands within progenitor spermatogonia. Importantly, this metabolic engagement occurs in concert with redox buffering and checkpoint activation, depicting an adaptive bioenergetic state rather than impending mitochondrial dysfunction. In this context, mitochondrial activity and genome surveillance appear co-optimized to preserve stem-cell integrity while enabling differentiation competence.

The molecular networks identified in this study also provide a framework for understanding how environmental and endocrine disruptors may compromise male fertility. A growing body of evidence links declining sperm quality and rising rates of male infertility to environmental exposures, including endocrine-disrupting chemicals, heavy metals, and dietary toxicants, many of which converge on oxidative stress pathways and kinase signalling cascades as common downstream mediators of damage^88^. In this context, toxicants that elevate testicular ROS or perturb key kinases such as PAK1, BUB1, or ABL2 would be predicted to disrupt precisely the antioxidant and genomic safeguarding networks defined here, thereby destabilizing spermatogonial integrity. Importantly, environmental perturbations of the germline extend beyond immediate fertility effects and may influence epigenetic programming across generations. Recent studies have demonstrated that environmental and metabolic exposures reshape the sperm epigenome, including alterations in histone modifications established during spermatogenesis, which influence placental development and set the trajectory for offspring health^89–92^. These findings raise the possibility that redox imbalance and kinase dysregulation within SSCs may not only impair spermatogenesis directly, but could also reprogram chromatin landscapes with intergenerational health impacts. Taken together, our dataset defines a series of kinase and antioxidant “nodes” that represent potential vulnerability points within the spermatogonial compartment. Future studies could leverage this framework to screen environmental chemicals against these defined phospho-signalling and redox pathways, enabling a more mechanistically informed approach to predicting reproductive toxicity. Such efforts may help bridge molecular-level perturbations in SSCs with broader epidemiological trends in male reproductive health.^93^

In conclusion, proteomic and phosphoproteomic data produced in this study have provided a platform of information with extensive translational potential. The identification of uniquely expressed / uniquely phosphorylated proteins and the activity of their associated pathways in SSCs provides a gateway for improved manipulation of these cells, for instance to improve SSC expansion and differentiation protocols for fertility restoration. The development of such stem cell technologies would provide transferable benefit from the paediatric oncofertility space through to wildlife conservation and livestock breeding programs^73^. Data presented within this manuscript also provide a resource of candidate biomarkers (e.g., PAK1/BUB1 activity, GSTT1 levels) that could be explored as a means to stratify idiopathic male-infertility patients. Ultimately, multifaceted approaches to understanding germ cell function (i.e. assessment at the transcript, protein, and phosphoprotein level) such as that presented in this manuscript hold substantial value in understanding the basis of male fertility across species, as well as the development of strategies to identify and treat infertility.

## Limitations of the study

This study utilised spermatogonia from postnatal day 6-8 mice because of the inherent enrichment of undifferentiated spermatogonia in the neonatal testis, facilitating isolation of more abundant populations than is possible from the adult testis, with less likelihood of contamination from other cell types. Regardless, there is evidence to suggest that spermatogonia from the adult testis may exhibit differences in their expression profile^48^ and metabolome^94^. Thus, a future direction would be to perform further analyses on spermatogonia from the adult testis, ideally integrating metabolomics with (single-cell) proteomics to capture cell-state heterogeneity and metabolic flux. Additionally, although pharmacological inhibitors were employed to infer functionality of selected kinases in undifferentiated spermatogonia in this study using primary cultures of undifferentiated spermatogonia, conditional knockout mouse lines for the kinase triad would need to be employed to validate causality *in vivo.* Correspondingly, CRISPR directed mutation at specific phosphosites would provide the ultimate confirmation of functional consequences associated with the changes in phosphorylation reported here. Finally, although knockout lines used in this study provided an informative snapshot of testis phenotype, statistical comparisons were not possible given that tissue from only two replicates per line is stored by the German Mouse Clinic. Thus, further investigation with additional replicates would be required to ascertain the statistical significance of the trends identified.

## ACKNOWLEDGEMENTS

Thank you to Nicole Cole from the University of Newcastle’s Central Analytical facilities for her assistance with FACS isolations. We thank Dr Ben Crossett and Angela Connolly from the Mass Spectrometry Core Facility at The University of Sydney, and the Academic and Research Computing Support team at University of Newcastle who provided High Performance Computing Infrastructure to support the bioinformatics analyses. We thank the technicians and animal caretakers of the German Mouse Clinic, and Robert E Braun and the IMPC working group for sharing their expertise in testis phenotyping.

## AUTHOR CONTRIBUTIONS

Conceptualization, D.A.S.B. and T.L; Methodology, D.A.S.B., A.S.M., F.D., P.S.B., J.M.O., R.T., V.G.D., H.F., M.H.A., C.C., and I.R.B.; Software, D.A.S.B.; Formal Analysis D.A.S.B., C.C., I.R.B., and T.L.; Investigation, D.A.S.B., A.S.M., F.D., P.S.B., C.C., I.R.B., and T.L.; Resources, D.A.S.B, B.N., V.G.D., H.F., M.H.A., and T.L.; Data Curation, D.A.S.B and T.L.; Validation, A.S.M., F.D., P.S.B., C.C., and I.R.B.; Writing – Original Draft D.A.S.B. and T.L.; Writing – Review & Editing D.A.S.B., B.N., A.S.M., F.D., P.S.B., R.T., V.G.D., H.F., M.H.A., J.M.O., C.C., I.R.B. and T.L.; Visualization, D.A.S.B and T.L.; Supervision, D.A.S.B, B.N., and T.L.; Funding Acquisition, T.L.;

## STAR METHODS

### KEY RESOURCES TABLE

| REAGENT or RESOURCE | SOURCE | IDENTIFIER |
| --- | --- | --- |
| **Antibodies** |  |  |
| Biotin conjugated CD90.2 (Thy-1.2), mouse monoclonal | Invitrogen | Cat#13-0902-82 |
| Anti-Biotin microbeads | Miltenyi | Cat#130-090-485 |
| Anti-Caspase 3, rabbit polyclonal | Abcam | Cat#ab13847 |
| Anti-cKIT, rat monoclonal, PE/Cy7 conjugated | Abcam | Cat#ab95676 |
| Anti-γH2AX (phospho S139), mouse monoclonal | Abcam | Cat#ab22551 |
| Anti-LIN28 (A177), rabbit polyclonal | Cell Signaling | Cat#3987S |
| Anti-STRA8, rabbit polyclonal | Abcam | Cat#ab49405 |
| Anti-SOX9, rabbit monoclonal | Abcam | Cat#ab185966 |
| Alexa Fluor anti-rabbit 594 | Thermo Fisher Scientific | Cat#A-11012 |
| Alexa Fluor anti-mouse 594 | Thermo Fisher Scientific | Cat# A-11005 |
| **Critical Commercial Assays** |  |  |
| Pierce™ BCA Protein Assay Kit | Thermo Fisher Scientific | Cat#23225 |
| Endopeptidase Lys-C | Wako Chemicals | Cat#129-02541 |
| Proteomics-grade modified trypsin | Sigma | Cat#T6567 |
| Titansphere TiO<sub>2</sub> Bulk Material | GL Sciences | Cat#5020-75000 |
| iClick™ TUNEL Andy Fluor™ 647 Imaging Assay | ABP Biosciences | Cat#A063 |
| FxCycle PI/Rnase Staining Solution | Invitrogen | Cat#F10797 |
| Live/dead Fixable far red dead cell stain | Invitrogen | Cat#L34973 |
| **Deposited Data** |  |  |
| Reference Proteomes – *Mus musculus* | UniProt | UP000000589;<br>RRID:SCR_004426 |
| Mouse spermatogonial stem cells and progenitor spermatogonia – RNA-seq | Helsel *et al*<sup>21</sup> |  |
| Mouse spermatogonial stem cells and progenitor spermatogonia - proteome | This paper | PRIDE: PXD066114<br>RRID:SCR_003411 |
| Mouse spermatogonial stem cells and progenitor spermatogonia - phosphoproteome | This paper | PXD066114<br>RRID:SCR_003411 |
| Mouse Genome Informatics | MGI<sup>95</sup> | RRID:SCR_006460 |
| The International Mouse Phenotyping Consortium | IMPC<sup>96</sup> | RRID:SCR_006158 |
| **Experimental Models / Organisms / Strains:** |  |  |
| Mouse: *Id4-eGfp* hemizygous | A gift from Professor Jon M. Oatley at Washington State University | N/A |
| Mouse: *Arcvf* homozygous knockout | German Mouse Clinic (GMC) / IMPC | Project # P02090 |
| Mouse: *Nucks1* homozygous knockout | GMC / IMPC | Project # P00908 |
| Mouse: *Rtn4* homozygous knockout | GMC / IMPC | Project # P01788 |
| Mouse: *Gstt1* homozygous knockout | GMC / IMPC | Project # P00886 |
| Mouse: *Epc2* heterozygous | GMC / IMPC | Project # P00876 |
| Mouse: *Cnot6l* homozygous knockout | GMC / IMPC | Project # P01808 |
| Mouse: *Mindy1* / *Fam63a* homozygous knockout | GMC / IMPC | Project # P00216 |
| **Other** |  |  |
| PAK1 Inhibitor (NVS-PAK1-1) | Selleck Chemicals | Cat #S8264 |
| BUB1 inhibitor (BAY-1816032) | Selleck Chemicals | Cat #S8945 |
| ABL2 inhibitor (Asciminib) | Selleck Chemicals | Cat #S8555 |
| All-trans retinoic acid | Stemcell technologies | Cat#72262 |
| **Software and Algorithms** |  |  |
| Proteome Discoverer 2.5 | Thermo Fisher Scientific | RRID:SCR_014477 |
| Perseus (version 2.0.7.0) | <sup>97</sup> | RRID:SCR_015753 |
| Ingenuity Pathway Analysis | Qiagen | RRID:SCR_008653 |
| GraphPad Prism (version 11.0.0 (93)) | Graph Pad | RRID:SCR_002798 |
| Biorender | Biorender.com | RRID:SCR_018361 |
| Image J | National Institute of Health | RRID: SCR_003070 |

### RESOURCE AVAILABILITY

#### Data and Code Availability

- The mass spectrometry proteomics data have been deposited to the ProteomeXchange Consortium (http://proteomecentral.proteomexchange.org) via the PRIDE partner repository^98^ with the dataset identifier PXD066114.

PRIDE Reviewer account details: Username; Password: oU2wxKqIhPbp

- All original code has been deposited at GitHub and is publicly available at https://github.com/DavidSBEire/ShinySpermagoniaCells as of the date of publication
- Any additional information required to reanalyse the data reported in this work paper is available from the Lead Contact upon request

#### Lead contact

Further information and requests for resources and reagents should be directed to and will be fulfilled by the Lead Contact, Tessa Lord.

#### Materials availability

Antibodies and reagents are available from the sources listed in the key resources table. Unless otherwise specified, all reagents were purchased from Merck.

### EXPERIMENTAL MODEL AND SUBJECT DETAILS

#### Ethics approval

All procedures involving animal use were approved by the University of Newcastle Animal Care and Ethics Committee (ACEC, approval numbers A-2019-907, A-2024-412 and A-2024-410). The *Id4-eGfp* mouse line^20^ was a gift from Dr Jon Oatley.

### METHOD DETAILS

#### Fluorescence activated cell sorting (FACS) and flow cytometry analysis

For cell sorting, single cell suspensions from *Id4-eGfp* testes were generated by trypsin-EDTA digestion as described previously^20^. In order to isolate sufficient protein for proteomic and phosphoproteomic analyses, testis cells were pooled for all males from a litter to create one biological replicate. Cell sorting (FACS) of ID4-eGFP^Bright^ (SSC) and ID4- eGFP^Dim^ (progenitor) populations of spermatogonia from P6-8 testes was conducted using a BD FACS AriaIIu using previously described gating parameters (^21^ and Figure 1A).

For flow cytometry analysis, assays (Key Resource Table) were performed as per the manufacturers instructions on a BD FACS Canto II.

#### Phosphoproteomic sample preparation of spermatogonia

All samples were prepared for phosphoproteomic analysis using methodology known as EasyPhos (EP)^19,22,99–101^. In brief, 250 µL of chilled lysis buffer [4% (w/v) sodium deoxycholate (SDC); 100 mM Tris-HCl (pH 8.5)], was added to each sample and immediately heated (95°C, 5 min) to inactivate endogenous proteases and phosphatases. Samples were sonicated (4 × 20 s cycles, 75% output power), and an aliquot taken to determine protein concentration using a bicinchoninic acid assay (BCA). All samples were diluted to equal protein amounts (80 µg) in 270 µL of lysis buffer into a 2 mL 96 deepwell plate. Samples were reduced and alkylated [100 mM Tris(2-carboxyethyl)phosphine hydrochloride; 400 mM 2-chloroacetemide], using a Thermomixer (Eppendorf; Hamburg, Germany), samples were incubated for 5 min at 45°C (1,500 rpm). Enzymatic digestion was achieved using Lys-C and trypsin, at an enzyme-to-substrate ratio of 1:100 (w/w) and incubated overnight at 37°C with shaking (1,500 rpm). To the digested peptides, 400µL of isopropanol (ISO) was added and mixed thoroughly for 30 s on the Thermomixer (1,500 rpm). Next, 100 µL of EP Enrichment buffer [48% (v/v) TFA / 8 mM potassium dihydrogen phosphate (KH2PO4)] was added to each sample and mixed for 30 s thoroughly (1,500 rpm). The plate containing the peptides was spun at 2,000 x g for 15 min (RT) and supernatants carefully transferred to a new clean 2 mL 96 deepwell plate. To each sample, resuspend in EP loading buffer [6% (v/v) TFA / 80% (v/v) ACN; concentration of mg / μl^−1^] 5mg of TiO_2_ beads were added and incubated at 40°C with shaking (2,000 rpm). Beads were pelleted (2,000 x g, 1 min) and the supernatant removed. The beads were washed five times with 1 mL EP wash buffer [5% (v/v) TFA / 60% (v/v) ISO], each mixed briefly (1,500 rpm) and supernatant removed after spinning (2,000 x g, 1 min). Beads were resuspended in 75 µL of EP transfer buffer [0.1% (v/v) TFA / 60% (vol/vol) ISO] and transferred to a C8 StageTip^102^. An additional 75 µL EP transfer buffer was added to each well to ensure all beads were retained and transferred to their C8 StageTip. To an in-house 3D printed 96-well StageTip centrifuge device^22,99^, each StageTip was secured and spun through to dryness (1,500 x g, 8 min). Next, 2 x 60 µL of EP elution buffer [200 μl of 25% ammonia solution / 800 μl of 40% (v/v) ACN.] was added to each StageTip and spun in PCR tubes (1,500 x g, 3 min). Immediately, the PCR tubes were lyophilized using a centrifuge concentrator (Eppendorf; Hamburg, Germany) at 45°C for 30 min, ensuring the samples were not completely dry (∼20 µL). To the partially dried samples, 100 µL of styrenedivinylbenzene-reverse phase sulfonated (SDB-RPS) loading buffer [1% (v/v) TFA in ISO] was added and each sample transferred to a SDB-RPS StageTip^102^ for desalting^99^. StageTips were centrifuged at 1,500 × g for 3 min and subsequently washed with 100µL 99% ethylacetate/1% TFA (1,500 × g, 3 min). Finally, StageTips were washed with 100 µL each of 99% ISO/1% TFA and 0.2% TFA/5% acetonitrile, eluted with 60 µL of 5% NH_4_OH/80% acetonitrile, dried by vacuum concentration and re-suspended in MS loading buffer (2% acetonitrile/0.3% TFA). Purified peptides were subjected to off-line high pH fractionation, prior to analysis by high resolution nano liquid chromatography tandem mass spectrometry (nLC-MS/MS), while phosphopeptides were subjected only to analysis by high resolution nLC-MS/MS.

#### nLC-MS/MS analysis

Reverse phase nLC-MS/MS was performed using an Eclipse Orbitrap MS coupled to a Dionex Ultimate 3000RSLC nanoflow high-performance liquid chromatography system (Thermo Fisher Scientific). Peptide separation was then achieved using an in-house packed column, SGE MyCapLC Kit (Kinesis) 300 mm x 150 mm, employing a 90 min gradient of acetonitrile for proteome samples (8 fractions per sample) and 120 minutes for phosphopeptides. Full MS/data dependent acquisition MS/MS mode was utilized on Xcalibur (Thermo Fisher Scientific; version 4.2.47), *for the proteome:* with the Orbitrap mass analyzer set at a resolution of 60,000, to acquire full MS with a range of 300 – 1400 m/z, incorporating an automatic gain control target of 4 x 10^5^ and maximum fill time of 60 ms, set in cycle time mode (3s). Multiply charged precursors ions were selected for higher-energy collision dissociation fragmentation with a normalized collisional energy of 30. MS/MS fragments were measured at an Orbitrap resolution of 15,000 using an automatic gain control target of 5 x 10^4^ and maximum fill times of 28 ms. *For phosphopeptides:* 60,000, to acquire full MS with a range of 300 – 1650 m/z, incorporating an automatic gain control target of 6 x 10^5^ and maximum fill time of 50 ms, set in cycle time mode (3s). Multiply charged precursors ions were selected for higher-energy collision dissociation fragmentation with a normalized collisional energy of 30. MS/MS fragments were measured at an Orbitrap resolution of 15,000 using an automatic gain control target of 1 x 10^5^ and maximum fill times of 120 ms.

#### Primary cultures of undifferentiated spermatogonia

Primary cultures were established using THY1+ undifferentiated spermatogonia isolated from P6-8 testes via magnetic activated cell sorting (MACS) using a THY1.2 biotin conjugated antibody (Invitrogen) and anti-biotin microbeads (Miltenyi), as per the manufacturer’s instructions. Cultures were maintained at 10% O_2_, 5% CO_2_ and 37°C, in mouse serum-fee medium (mSFM) supplemented with glial-derived neurotrophic factor (GDNF, 20 ng/ml) (R&D systems, MN, USA) and fibroblast growth factor (FGF2, 1 ng/ml) (Sigma Aldrich)^103^. Cultures were seeded on a feeder layer of mitotically inactivated SNL76/7 mouse embryonic fibroblasts (ATCC, VA, USA) and were passaged every 6-8 days onto fresh feeders. Media was replaced every second day.

For kinase inhibitor studies, spermatogonia were dislodged from feeder cells using gentle pipetting, subjected to trypsin digestion to create a single cell suspension, and re- plated in feeder free conditions. Cells were then incubated with the desired concentration of inhibitor (Key Resource Table) or vehicle (DMSO) for 24 h prior to analysis.

#### Apoptosis assays: TUNEL and Caspase 3

In order to identify apoptosis in spermatogonia treated with kinase inhibitors, cells were fixed in 4% paraformaldehyde for 30 min and settled on poly-L-lysine coated coverslips. Cells were permeabilised in 0.2% Triton-X (Sigma Aldrich) in PBS for 30 min at room temperature. As a positive control for the TUNEL assay (ABP Biosciences) a subset of cells were treated with DNase I, and all samples were treated with TUNEL reaction cocktail and iClick reaction cocktail as per the manufacturer’s instructions.

For Caspase 3 detection, coverslips were incubated in blocking solution (3% BSA/PBS) then incubated in primary antibody diluted 1/200 in 1% BSA/PBS and incubated overnight at 4°C. The following day, cells were incubated in secondary antibody (Alexa Fluor 594) at a concentration of 1/1000 for 2 h at room temperature. Cells were counterstained with DAPI (1:1000 for 5 mins) and coverslips were mounted on slides using Mowiol. Coverslips were imaged on a Zeiss Axio A.2 fluorescence microscope or an FV1000 confocal microscope (Olympus, Tokyo, Japan). For TUNEL staining, at least 100 cells were counted per treatment / per replicate to establish the percentage of TUNEL-positive cells in the population. For Caspase 3, intensity of staining was quantified using Corrected Total Cell Fluorescence (CTCF) values calculated using Image J (National Institute of Health). Again, at least 100 cells per treatment / per replicate were assessed.

#### Detection of DNA damage

In order to detect DNA damage (double strand breaks) in spermatogonia in response to kinase inhibitor treatments, an immunocytochemistry protocol was adopted using a γH2AX antibody (Key Resource Table). Cells were prepared and mounted on coverslips as described above, with the anti-γH2AX primary antibody being diluted 1/500 in 1% BSA/PBS and incubated with cells overnight at 4°C. Secondary antibody and mounting steps were conducted as above. For each replicate, at least one hundred cells were counted per treatment to establish the percentage of γH2AX-positive cells (those with bright fluorescent signal).

#### Phenotyping of knockout mouse lines

Phenotyping and access to all knockout mouse lines used in this study (Table S7) was via the German Mouse Clinic and The International Mouse Phenotyping Consortium (IMPC)^70,84,104^. Mouse models were generated using the IMPC targeting strategy with CRISPR/Cas9 or ES cell technology at Helmholtz Munich (https://www.mousephenotype.org/understand/the-data/allele-design/)^68,69^. After genotyping, heterozygous × heterozygous matings were set up to generate sufficient mutant mice with littermate +/+ controls for phenotyping analysis at the German Mouse Clinic as described^68^, and in agreement with the standardized phenotyping pipeline of the IMPC including histopathological analysis (https://www.mousephenotype.org/impress/PipelineInfo?id=14).

Testes and epididymides from control and mutant mice (16 weeks of age, n = 2 per genotype) were fixed in neutral buffered formalin, processed and embedded in paraffin. For each testis, two regions separated by 100 μm, were serially sectioned (3 µm thick) at both ends. The first section from each region (#1 and #33) was stained with hematoxylin and eosin (H&E) to assess testis and epididymis morphology. The adjacent sections (#2 and #34) were immunostained for SOX9 to quantify atrophic seminiferous tubules, defined as tubules containing only SOX9-positive Sertoli cells. Subsequent sections from the first region (#3 and #4) were immunostained for LIN28A and STRA8 to identify undifferentiated and differentiating spermatogonia, respectively. Immunohistochemistry was performed on a BOND RX^m^ automated stainer (Leica Biosystems, Germany). Briefly, sections were deparaffinized and antigen retrieval was carried out for 30 minutes using either citrate buffer (BOND Epitope Retrieval Solution 1 – H1: SOX9) or ethylenediaminetetraacetic acid (EDTA) (Bond Epitope Retrieval Solution 2 – H2: LIN28A, STRA8). After a 30 minute blocking step, sections were incubated for 60 minutes with primary antibodies against SOX9 (1:1200), LIN28A (1:800) or STRA8 (1:4000) (Key Resources Table). Anti-rabbit secondary antibodies were applied for 30 minutes and detection was performed using BOND Polymer Refine Detection DAB (3,3’-diaminobenzidine). Sections were counterstained with hematoxylin. Negative controls were processed identically except that primary antibody was omitted and replaced with antibody diluent. All slides were scanned with a NanoZoomer S60 slide scanner (Hamamatsu, Japan) and evaluated using NDP.View2 Plus Software. The histological analyses were performed blind by at least two pathologists.

For image analysis, H&E-stained testis and epididymis images were analysed from each of the 42 knockout mouse lines and annotated according to a protocol developed by the IMPC Infertility working group. Histological annotations were recorded using an Excel database with standard adult mouse anatomy (MA) and mouse pathology (MPATH) ontologies^105–107^. Within this framework, MPATH terms (listed in Table S7) were assigned together with a severity score to provide a semi-quantitative assessment of histological lesions. A score of 0 indicated the absence of detectable abnormalities and scores 1, 2, 3, 4 and 5 corresponded to mild, moderate, marked, marked-to-severe and severe changes, respectively.

In seven knockout lines (*Gstt1, Epc2, Mindy1/Fam63a, Cnot6l, Nucks1, Rtn4*, and *Arvcf*) assigned with the MPATH terms MPATH:87 (Germ Cell Defect) and MPATH:797 (Spermatogenesis Defect), tubular atrophy was further quantified using H&E and SOX9 immunohistochemistry by determining the number of seminiferous tubules containing only SOX9-positive Sertoli cells (Sertoli cell-only tubules)^108^. In adult mice, the presence of a small number (1-5) of Sertoli-only tubules is generally considered a spontaneous background finding. In contrast, the presence of 6 or more such tubules distributed across multiple areas of the testis, particularly when accompanied by reduced epididymal sperm content, is considered indicative of testicular atrophy^71^. In this assessment, two sections at a distance of 100 µm (#1, #33) were assessed for SOX9-Sertoli cell only-tubules.

For evaluation of undifferentiated and differentiating spermatogonia, LIN28A and STRA8 immunostainings were quantified, respectively. The number of LIN28A-positive cells was determined in seminiferous tubule cross sections within a 6 mm^2^ area of testicular tissue from the left and right testes. The proportion of STRA8-positive tubule cross sections was calculated by examining a minimum of 60 tubule cross sections from each left and right testis section. For the *Mindy1*/*Fam63a* mouse line, prolonged formalin fixation compromised antigen preservation, preventing adequate epitope retrieval and consequently adequate immunohistochemical detection of the proteins.

### QUANTIFICATION AND STATISTICAL ANALYSIS

#### Proteomic data processing and analysis

Consistent with previous studies^99,100,109–113^, database searching of raw files were performed separately using Proteome Discoverer 2.5 (Thermo Fisher Scientific). SEQUEST HT was used to search against the UniProt *Mus musculus* database (25,444 sequences, downloaded 29^th^ November 2022). Database searching parameters included up to two missed cleavages, a precursor mass tolerance set to 10 ppm and fragment mass tolerance of 0.02 Da. Trypsin was designated as the digestion enzyme. Cysteine carbamidomethylation was set as a fixed modification while phosphorylation (S, T, Y) was designated as a dynamic modification. Interrogation of the corresponding reversed database was also performed to evaluate the false discovery rate (FDR) of peptide identification using Percolator on the basis of q-values, which were estimated from the target-decoy search approach. To filter out target peptide spectrum matches over the decoy-peptide spectrum matches, a fixed FDR of 1% was set at the peptide level. Label-free quantification was performed using Proteome Discoverer nodes “Minora Feature Detector”, “Feature Mapper”, and “Precursor Ions Quantifier” nodes as described previously^113^. Fold change and significance (*t*-test) comparative testing between sample groups was carried out within the Proteome Discoverer 2.5 suite. The peptide list was exported from Proteome Discoverer 2.5 as an Excel file and further refined to include only those with a quantitative value in all replicates of at least one of the sample groups. The refined peptide list was loaded into Perseus, version 2.0.7.0^114^, for the generation of scatter plots, principal component analysis, and heatmaps. Basic data handling, if not otherwise stated, was conducted using Microsoft Excel 365 (Version 16.98, Microsoft Corporation, Redmond, WA) and GraphPad Prism version 11.0.0(93) for Windows (GraphPad Software; San Diego, CA).

#### Ingenuity pathway analysis

For each of the sample groups, where a phosphopeptide or protein was expressed in all replicates, the corresponding UniProt accession numbers and transformed ratios were analysed using Ingenuity Pathway Analysis software (IPA®, Qiagen) as previously described^19,99,112,113,115,116^. SSCs and progenitor proteomes (8,450 / 8,464; 99.8%) were firstly analysed on the basis of predicted subcellular location and classification (other excluded). Further analyses focused on all significantly altered proteins and phosphopeptides (*p-*value ≤ 0.05) between SSCs and progenitor populations. Importantly, the IPA phosphorylation specific knowledge base was utilized for phosphoproteome data, ensuring the biological predictions are based upon patterns of phosphorylation changes and not protein expression. Canonical pathway, upstream regulators, and disease and function analyses were assessed using; *p*-value, an enrichment measurement of the overlapping proteins from the dataset in a particular pathway, function or regulator^117^ and; *Z*-score, a prediction scoring system of activation or inhibition based upon statistically significant patterns in the dataset and prior biological knowledge manually curated in the Ingenuity Knowledge Base. To elucidate the most significant changes in our analyses, we applied a stringency criteria of -log10 *p*-value of ≥ 1.3, and a Z-score of (inhibition) -2 ≤ Z ≥ 2 (activation) in at least one group^99,112,113,118^. For disease and function assessment we restricted the analysis to ‘molecular and cellular functions’ and ‘physiological system development and function’.

#### Kinase identification

In addition to the direct identification of kinases within the phosphoproteomic dataset and the upstream regulator feature of IPA, the dataset was subjected to a kinase-substrate analysis utilizing the PhosphoSitePlus database (accessed 23^rd^ April 2024, http://www.phosphosite.org/), an open-source curated resource for investigating the importance of experimentally observed post-translational modifications in the regulation of biological processes^62^, as previously described^19^. KinMap^119^ was utilized in the visual representation of the findings.

#### Shiny Application development

In accordance with Shiny blueprint outlined by the ShinySperm Application family^61,92,120,121^, web-based applications support the accessibility and interpretability of datasets allowing for effective data-driven insights by the field. The full coding script supporting ShinySpermatogoniaCells (https://reproproteomics.shinyapps.io/ShinySpermatogoniaCells/), can be downloaded from GitHub: https://github.com/DavidSBEire/ShinySpermagoniaCells. In brief, the ShinySpermatogoniaCells application was built using the shiny package (version 1.9.1) on RStudio (version 2024.04.1+748), with base *R* (version 4.3.3, 2024-02-29). Supporting the functionality and aesthetics of this application are several packages, including: DT, dplyr, ggplot2, openxlsx, plotly, readxl, and shinydashboard.

#### Statistical Analysis

Phosphoproteomic analyses were performed using SSC and progenitor populations isolated from an ID4-eGFP mouse line, as described above (n = 4 biological replicates). Differentially accumulated or phosphorylated spermatogonia proteins were defined as those with a fold-change ± 1.5 and *p-*value ≤ 0.05. All other data were assessed for normality using a Shapiro-Wilk normality test. Normally distributed data were analysed via Student’s t-tests to detect differences between treatment groups. Data not normally distributed were analysed by a Mann-Whitney test. Differences between groups were considered significant when *p* ≤ 0.05. The number of biological replicates used in each experiment are presented in figure captions. Graphical data were prepared using GraphPad Prism (version 11.0.0 (93)) and are presented as mean values ± SEM.

## SUPPLEMENTAL FIGURE LEGENDS

**Figure S1:** Dose response analyses for dose selection of the following kinase inhibitors: PAK1 Inhibitor (NVS-PAK1-1), BUB1 inhibitor (BAY-1816032), ABL2 inhibitor (Asciminib) (relates to Figure 4). All incubations were conducted over 24 hours using primary cultures of undifferentiated spermatogonia from ID4-eGFP mice.

**A)** Total cell number normalised to the vehicle control (DMSO). The IC50 for each inhibitor is listed above each histogram. Histograms depict mean ± SEM for n=4 independent biological replicates.

**B)** Percentage of viable cells (Live/Dead stain) following inhibition of selected kinases as compared to the vehicle control. Histograms depict mean ± SEM for n=4 independent biological replicates.

**C)** Mean fluorescence intensity (MFI) of the ID4-eGFP transgene (a reflection of SSC content) following inhibition of selected kinases as compared to the vehicle control. Histograms depict mean ± SEM for n=4 independent biological replicates.

**Figure S2:** Representative images accompanying data in Figure 4E, F, G, depicting apoptosis and DNA damage in spermatogonia following inhibition of selected kinases.

**A)** TUNEL staining in undifferentiated spermatogonia following treatment of primary cultures with kinase inhibitors for PAK1, BUB1, ABL2, or Trio, as compared to a vehicle (DMSO) treated control. Negative control had TUNEL enzyme omitted, and positive control was treated with DNase. Blue staining is DAPI, purple staining is TUNEL. Scale bar = 20 µm.

**B)** Cleaved Caspase 3 in undifferentiated spermatogonia following treatment of primary cultures with kinase inhibitors for PAK1, BUB1, ABL2, or Trio, as compared to a vehicle (DMSO) treated control. Negative control had primary antibody omitted. Blue staining is DAPI, red staining is Caspase 3. Scale bar = 20 µm.

**C)** ãH2AX in undifferentiated spermatogonia following treatment of primary cultures with kinase inhibitors for PAK1, BUB1, ABL2, or Trio, as compared to a vehicle (DMSO) treated control. Negative control had primary antibody omitted. Blue staining is DAPI, red staining is γH2AX. Scale bar = 20 µm.

**Figure S3:** Photomicrographs of H&E (hematoxylin and eosin)-stained testes and caudal epididymides (mice 1 and 2) from 16-week-old wild-type (WT), *Cnot6l-/-, Epc2+/-, Fam63a-/-, Gstt1-/-, Arvcf+/-, Nucks1-/-,* and *Rtn4-/-* mice. Insets show higher magnification views of selected seminiferous tubules and caudal epididymal regions. Images accompany data presented in Figure 5 and Table S7. Scale bar = 500 µm and 50 µm in magnified inset.

**Figure S4:** Assessment of undifferentiated and differentiating spermatogonia in knockout mouse lines that had elevated numbers of atrophic tubules (accompanying data in Figure 5 and S3).

**A)** Comparison of the number of LIN28+ undifferentiated spermatogonia per tubule across 4 knockout mouse lines as compared to age matched wild type controls. Histograms depict mean ± SEM. Knockout lines are n = 2 biological replicates and n = 6-8 technical replicates (individual testis sections). For wild type comparison, analysis was performed across n = 4 biological replicates and n = 16 technical replicates.

**B)** Comparison of the number of tubules in stage VII-VIII of the seminiferous epithelium (i.e. containing STRA8+ spermatogonia) across 4 knockout mouse lines as compared to age matched controls. Histograms depict mean ± SEM. Knockout lines are n = 2 biological replicates and n = 6-8 technical replicates (individual testis sections). For wild type comparison, analysis was performed across n = 4 biological replicates and n = 16 technical replicates.

**C)** Immunohistochemical detection of LIN28A proteins in testes from 16-week-old wild-type (WT) control and Gstt1-/-, Cnot6l-/-, Nucks1-/- and Rtn4-/- mice. Insets show higher magnification views of cross-sections of seminiferous tubules.

**D)** Immunohistochemical detection of STRA8 proteins in testes from 16-week-old wild-type (WT) control and Gstt1-/-, Cnot6l-/-, Nucks1-/- and Rtn4-/- mice. Insets show higher magnification views of cross-sections of seminiferous tubules.

## SUPPLEMENTAL TABLES

**Table S1** Proteomic characterisation of SSC and progenitor spermatogonia. Data related to Figures 1 and 2.

**Table S2** Comparison of RNA-seq and proteome of SSC and progenitor populations. Data related to Figure 2.

**Table S3** Ingenuity Pathway analysis outputs related to significant changes in the proteome between SSC and progenitor populations. Data related to Figure 2.

**Table S4** Phosphoproteomic characterisation of SSC and progenitor spermatogonia. Data related to Figure 3.

**Table S5** Ingenuity Pathway analysis outputs related to significant changes in the phosphoproteome between SSC and progenitor populations. Data related to Figure 3.

**Table S6** Full list of kinases mapped through proteomic and phosphoproteomics analysis, IPA and PhosphositePlus mapping. Data related to Figure 3.

**Table S7** Characterisation of knockout mouse line phenotypes. Data related to Figure 5.

