## Supplemental data for "A proteomic and phosphoproteomic comparison of mouse spermatogonial stem cells and progenitor spermatogonia"

### SUPPLEMENTAL FIGURE 1

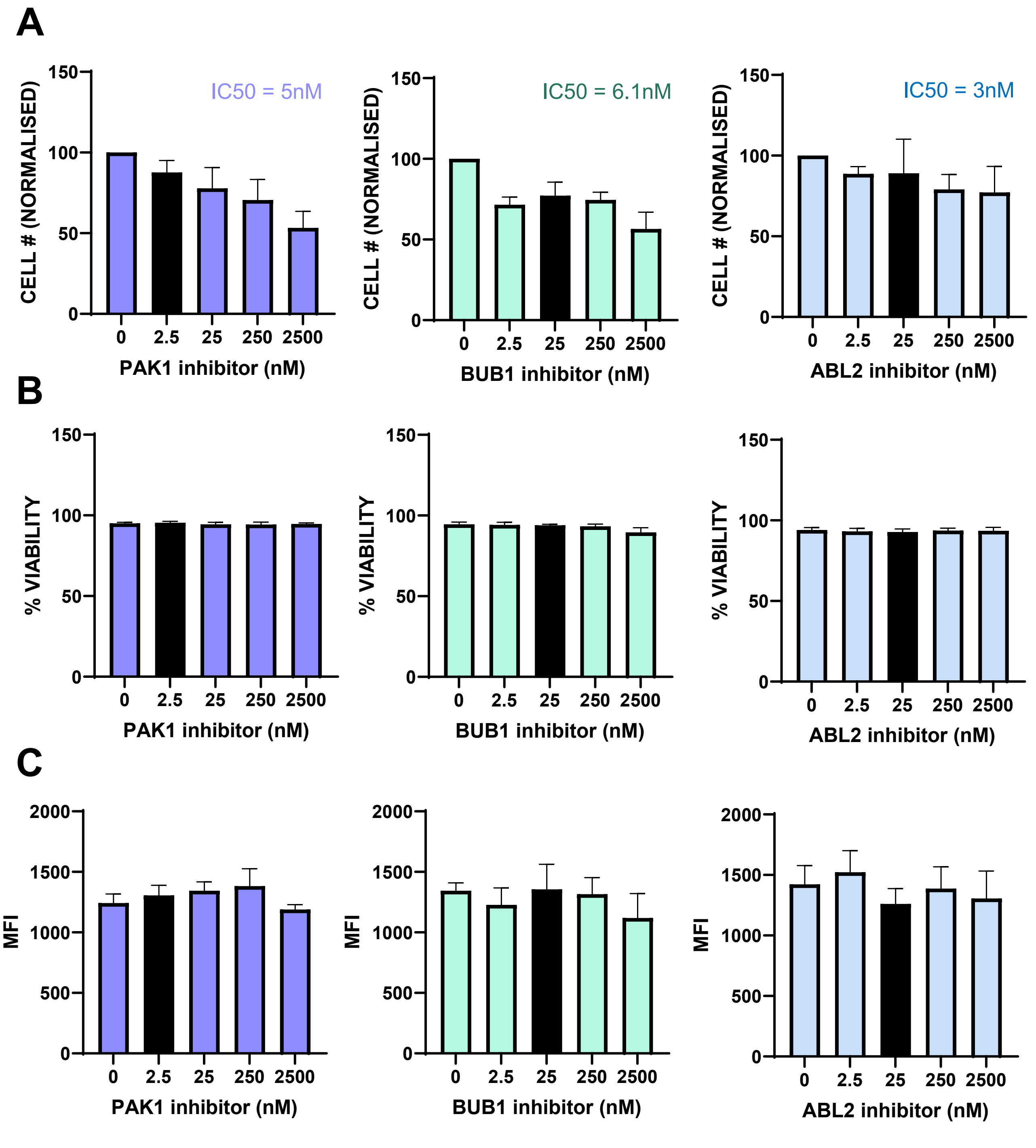


**Figure S1:** Dose response analyses for dose selection of the following kinase inhibitors: PAK1 Inhibitor (NVS-PAK1-1), BUB1 inhibitor (BAY-1816032), ABL2 inhibitor (Asciminib) (relates to Figure 4). All incubations were conducted over 24 hours using primary cultures of undifferentiated spermatogonia from ID4-eGFP mice.

**A)** Total cell number normalised to the vehicle control (DMSO). The IC50 for each inhibitor is listed above each histogram. Histograms depict mean ± SEM for n=4 independent biological replicates.

**B)** Percentage of viable cells (Live/Dead stain) following inhibition of selected kinases as compared to the vehicle control. Histograms depict mean ± SEM for n=4 independent biological replicates.

**C)** Mean fluorescence intensity (MFI) of the ID4-eGFP transgene (a reflection of SSC content) following inhibition of selected kinases as compared to the vehicle control. Histograms depict mean ± SEM for n=4 independent biological replicates.

### SUPPLEMENTAL FIGURE 2

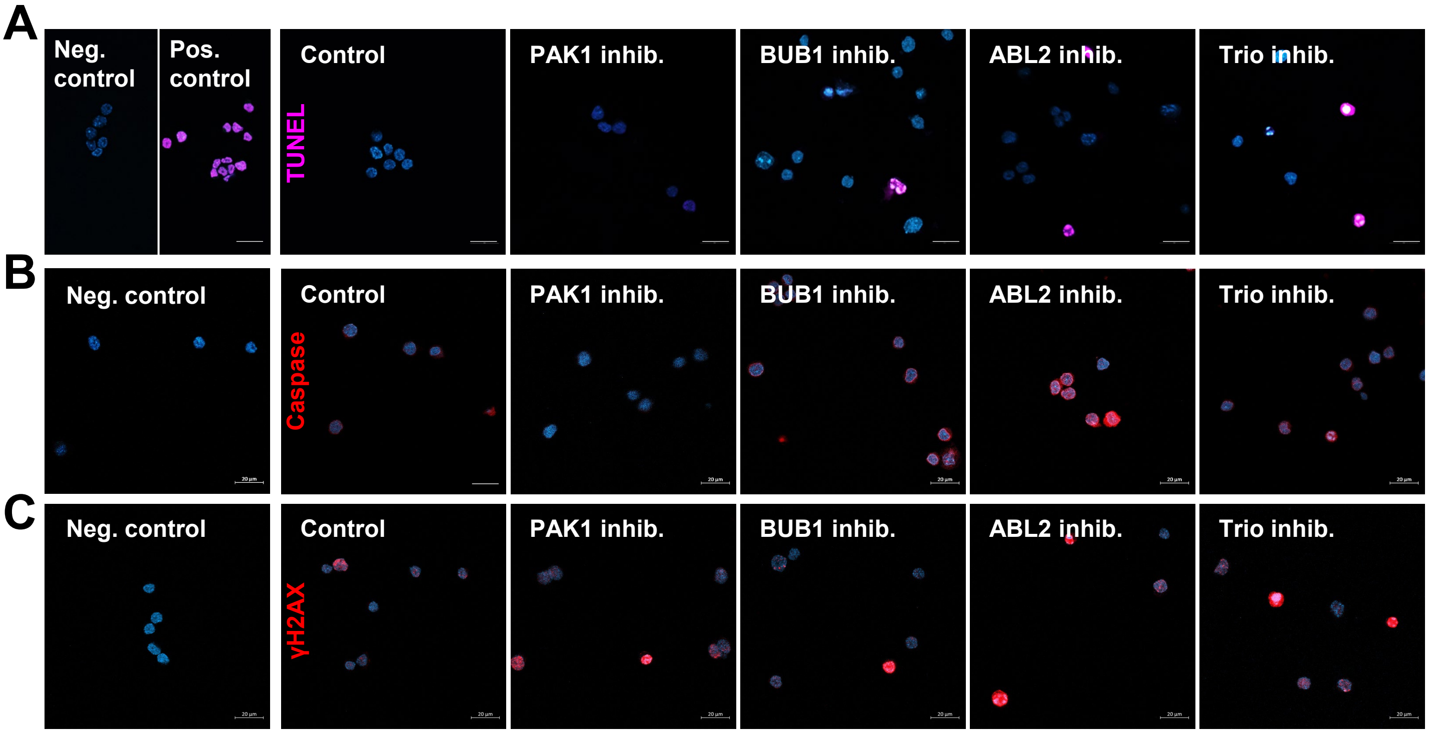


**Figure S2:** Representative images accompanying data in Figure 4E, F, G, depicting apoptosis and DNA damage in spermatogonia following inhibition of selected kinases.

**A)** TUNEL staining in undifferentiated spermatogonia following treatment of primary cultures with kinase inhibitors for PAK1, BUB1, ABL2, or Trio, as compared to a vehicle (DMSO) treated control. Negative control had TUNEL enzyme omitted, and positive control was treated with DNase. Blue staining is DAPI, purple staining is TUNEL. Scale bar = 20 µm.

**B)** Cleaved Caspase 3 in undifferentiated spermatogonia following treatment of primary cultures with kinase inhibitors for PAK1, BUB1, ABL2, or Trio, as compared to a vehicle (DMSO) treated control. Negative control had primary antibody omitted. Blue staining is DAPI, red staining is Caspase 3. Scale bar = 20 µm.

**C)** γH2AX in undifferentiated spermatogonia following treatment of primary cultures with kinase inhibitors for PAK1, BUB1, ABL2, or Trio, as compared to a vehicle (DMSO) treated control. Negative control had primary antibody omitted. Blue staining is DAPI, red staining is γH2AX. Scale bar = 20 µm.

### SUPPLEMENTAL FIGURE 3

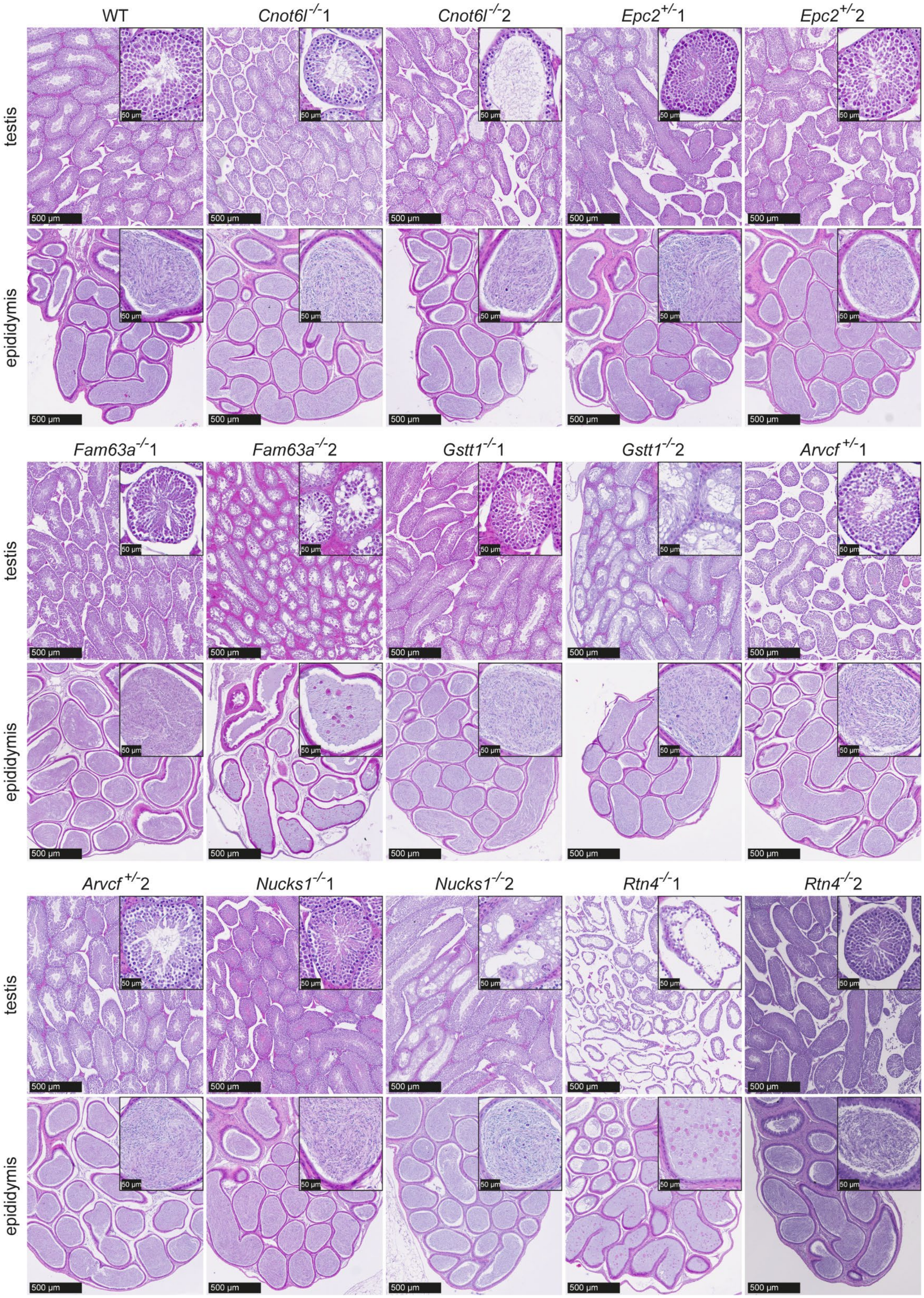


**Figure S3:** Photomicrographs of H&E (hematoxylin and eosin)-stained testes and caudal epididymides (mice 1 and 2) from 16-week-old wild-type (WT), *Cnot6l-/-, Epc2+/-, Fam63a-/-, Gstt1-/-, Arvcf+/-, Nucks1-/-,* and *Rtn4-/-* mice. Insets show higher magnification views of selected seminiferous tubules and caudal epididymal regions. Images accompany data presented in Figure 5 and Table S7. Scale bar = 500 μm and 50 μm in magnified inset.

### SUPPLEMENTAL FIGURE 4

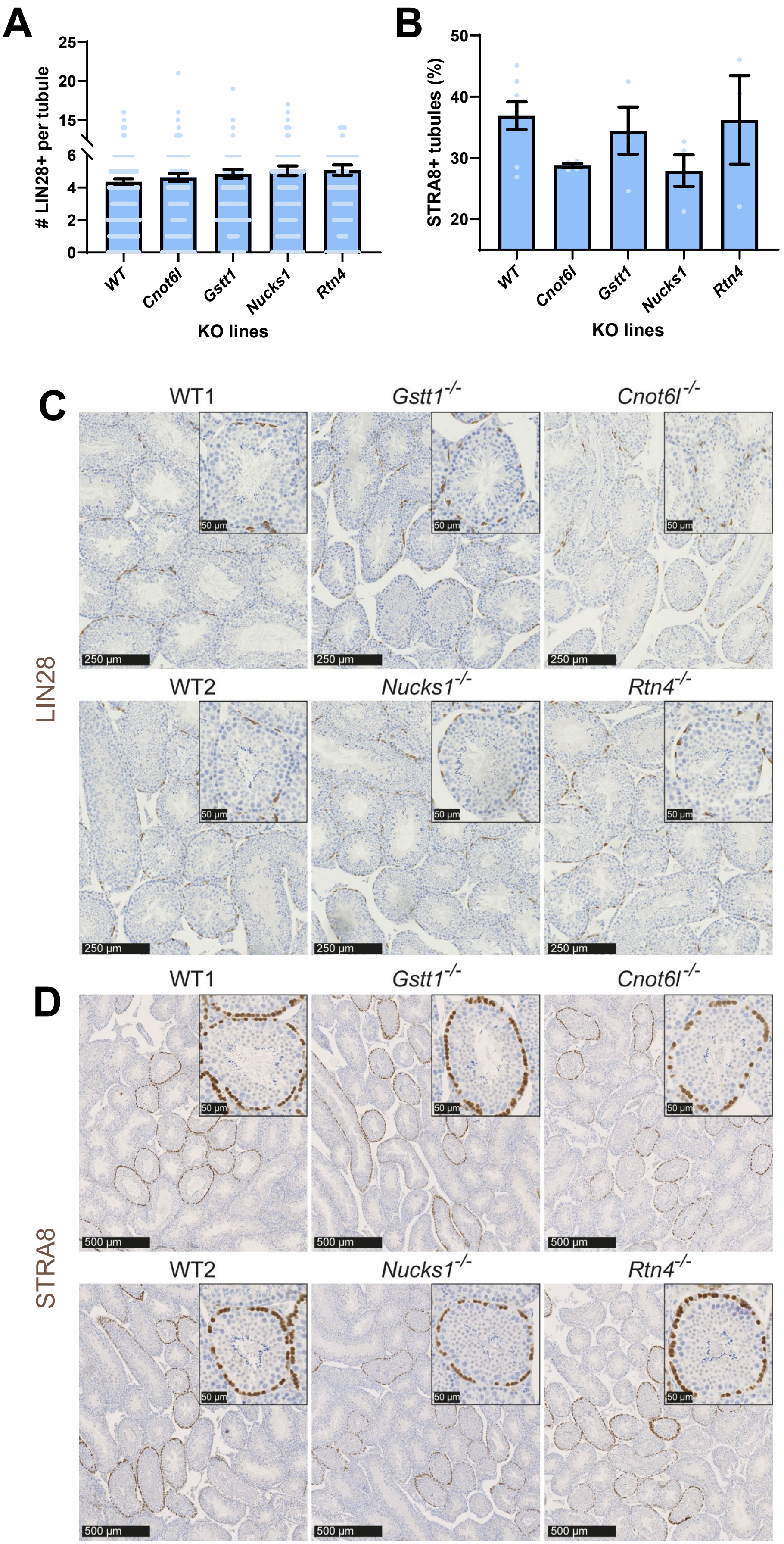


***Figure legend next page…***

**Figure S4:** Assessment of undifferentiated and differentiating spermatogonia in knockout mouse lines that had elevated numbers of atrophic tubules (accompanying data in Figure 5 and S3).

**A)** Comparison of the number of LIN28+ undifferentiated spermatogonia per tubule across 4 knockout mouse lines as compared to age matched wild type controls. Histograms depict mean ± SEM. Knockout lines are n = 2 biological replicates and n = 6-8 technical replicates (individual testis sections). For wild type comparison, analysis was performed across n = 4 biological replicates and n = 16 technical replicates.

**B)** Comparison of the number of tubules in stage VII-VIII of the seminiferous epithelium (i.e. containing STRA8+ spermatogonia) across 4 knockout mouse lines as compared to age matched controls. Histograms depict mean ± SEM. Knockout lines are n = 2 biological replicates and n = 6-8 technical replicates (individual testis sections). For wild type comparison, analysis was performed across n = 4 biological replicates and n = 16 technical replicates.

**C)** Immunohistochemical detection of LIN28A proteins in testes from 16-week-old wild-type (WT) control and Gstt1-/-, Cnot6l-/-, Nucks1-/- and Rtn4-/- mice. Insets show higher magnification views of cross-sections of seminiferous tubules.

**D)** Immunohistochemical detection of STRA8 proteins in testes from 16-week-old wild-type (WT) control and Gstt1-/-, Cnot6l-/-, Nucks1-/- and Rtn4-/- mice. Insets show higher magnification views of cross-sections of seminiferous tubules.
